# Replication intermediate architecture reveals the chronology of DNA damage tolerance pathways at UV-stalled replication forks in human cells

**DOI:** 10.1101/2020.10.12.336107

**Authors:** Yann Benureau, Caroline Pouvelle, Eliana Moreira Tavares, Pauline Dupaigne, Emmanuelle Despras, Eric Le Cam, Patricia Kannouche

**Author notes:** co-last authors (Emmanuelle Despras, Eric Le Cam, Patricia Kannouche).

## Abstract

DNA lesions in S phase threaten genome stability. The DNA damage tolerance (DDT) pathways overcome these obstacles and allow completion of DNA synthesis by the use of specialised translesion (TLS) DNA polymerases or through recombination-related processes. However, how these mechanisms coordinate with each other and with bulk replication remain elusive. To address these issues, we monitored the variation of replication intermediate architecture in response to ultraviolet irradiation using transmission electron microscopy. We show that the TLS polymerase *η*, able to accurately bypass the major UV lesion and mutated in the skin cancer-prone xeroderma pigmentosum variant (XPV) syndrome, acts at the replication fork to resolve uncoupling and prevent post-replicative gap accumulation. Repriming occurs as a compensatory mechanism when this on-the-fly mechanism cannot operate, and is therefore predominant in XPV cells. Interestingly, our data support a recombination-independent function of RAD51 at the replication fork to sustain repriming. Finally, we provide evidence for the post-replicative commitment of recombination in gap repair and for pioneering observations of *in vivo* recombination intermediates. Altogether, we propose a chronology of UV damage tolerance in human cells that highlights the key role of pol*η* in shaping this response and ensuring the continuity of DNA synthesis.

## Introduction

Accurate DNA replication is necessary to maintain genome stability. However, various endogenous or exogenous DNA damage constantly threaten the progression of the high-fidelity replicative DNA polymerases. The DNA damage response (DDR) organizes the multiple intricate molecular mechanisms controlling cell cycle progression, DNA repair and DNA damage tolerance (DDT). DDT comprises two pathways, conserved throughout evolution, that allow completion of DNA replication in spite of the presence of DNA lesions in the parental strands. Translesion synthesis (TLS) relies on the replacement of stalled replicative DNA polymerases by specific error-prone specialised DNA polymerases able to directly insert nucleotides in front of the lesion, in an error-free or an error-prone manner^1^. Alternatively, homologous recombination (HR)-like mechanisms, globally referred as damage avoidance (DA) or homology-dependent repair (HDR), use the sister chromatid to circumvent the damage by strand exchange or template switch^2^.

Whether the DDT pathways compete, compensate or collaborate is still unclear^3^. Data from bacteria and yeast suggest a chronological occurrence, with TLS acting before HDR^4, 5^. In human, a certain specialisation of the TLS polymerases is illustrated by the *xeroderma pigmentosum* variant (XPV) syndrome, a hereditary skin cancer-prone disease caused by inactivating mutations in the gene coding for the Y-family TLS polymerase *η* (pol*η*). Pol*η* has the unique capacity to accurately overcome cyclobutane thymine dimers (TT-CPDs), which are the most abundant lesions induced by UV irradiation^6–8^. Error-prone bypass by alternative TLS polymerases accounts for the hypermutability to UV of XPV cells^9–12^. XPV cells also show increased recombination, as evidenced by a higher number of sister chromatid exchanges (SCEs)^13^, but it is currently unclear whether this contributes to the disease. Hence, pol*η* plays a unique role in the tolerance of UV damage and cannot be efficiently and accurately replaced by other DDT mechanisms.

How DDT coordinates with replication fork (RF) progression is also a matter of debate and DDT timing was proposed to impact on its accuracy and on the correct maintenance of the epigenetic marks in the surrounding chromatin^14, 15^. Various studies showed that TLS and HR are post-replicative mechanisms in yeast, proceeding uncoupled from the bulk replication to fill in gaps left behind RFs through stalling of the synthesis of an Okazaki fragment on the lagging strand or repriming of a stalled leading strand^5, 16–18^. A similar post-replicative model was proposed for TLS in human cells^14, 19^ and the recently identified primase-TLS polymerase PrimPol has been proposed to mediate repriming at stalled RFs^20–23^. However, numerous studies showed that TLS contributes to RF progression^12, 20, 24–27^ and it is likely that TLS can take place both at and behind the RFs in vertebrates^25, 28, 29^. ssDNA gaps are excellent substrates for HR in yeast^30, 31^. Adar and colleagues showed, in a plasmid model, that ssDNA gaps located opposite to lesions can be filled in by RAD51-dependent HR mechanism in mammalian cells^32^. Yet, several recombination-like RF restart pathways were also described. Moreover, in the last few years, fork reversal became a widespread structural model to describe the response of RFs to any kind of replication stress, from nucleotide pool imbalance to bulky DNA lesions^33, 34^. RAD51 plays a critical role in this process, although its precise functions in RF remodelling, protection from nucleases and restart is not entirely clear.

In this work, we investigated the coordination of genome replication with DDT pathways in human fibroblasts following exposure to UVC. We used transmission electron microscopy (TEM) to assess the variations of replication intermediate (RI) architecture at early and late time points after irradiation in pol*η*-proficient and pol*η*-deficient backgrounds, coupled to analysis of RF progression by DNA fiber assay and monitoring of the recruitment of specific proteins to replicating chromatin by iPOND. We showed that pol*η*activity configures the immediate response of RFs to UV-induced lesions and promotes their progression and the continuity of DNA synthesis. Its absence diverts lesion tolerance at the RF to post-replicative mechanisms and we present evidence that PrimPol contributes to but is not the main actor of repriming in XPV cells. Finally, our observations suggest that recombination intermediates appear later after UV and correlate with the accumulation of RAD51 behind RFs and its requirement for gap repair. Interestingly, pol*η* deficiency also led to the recruitment of RAD51 at the RFs where it helps to overcome RF uncoupling in a strand invasion-independent manner, presumably by contributing to repriming events.

## Results

### Polη is recruited in the vicinity of replication forks to promote their progression after UVC irradiation

It is well documented that pol*η* accumulates at replication foci after UV in human cells^35, 36^. However, it remains unclear if this recruitment occurs at stalled RFs or during post-replicative gap filling. We therefore performed iPOND, a procedure that allows discriminating between proteins enriched at and behind RFs^37^. Cells were pulse-labelled with 5-ethynyl-2’-deoxyuridine (EdU) after UVC irradiation and cross-linked immediately or after a thymidine chase. Proteins bound to EdU-labelled DNA were pulled-down and analysed by western blot (Figure 1a). Pol*η* could readily be detected in the mock-irradiated sample, as we have previously shown^38^. Pol*η* amounts increased after UV and, like PCNA amounts, decreased with chase, showing that pol*η* is recruited preferentially in the vicinity of RFs after UV. Interestingly, RAD51 showed a mirrored dynamic, with levels increasing in post-replicative chromatin, which suggests a spatio-temporal separation between pol*η*-dependent TLS and recombination events.

**Figure 1.**
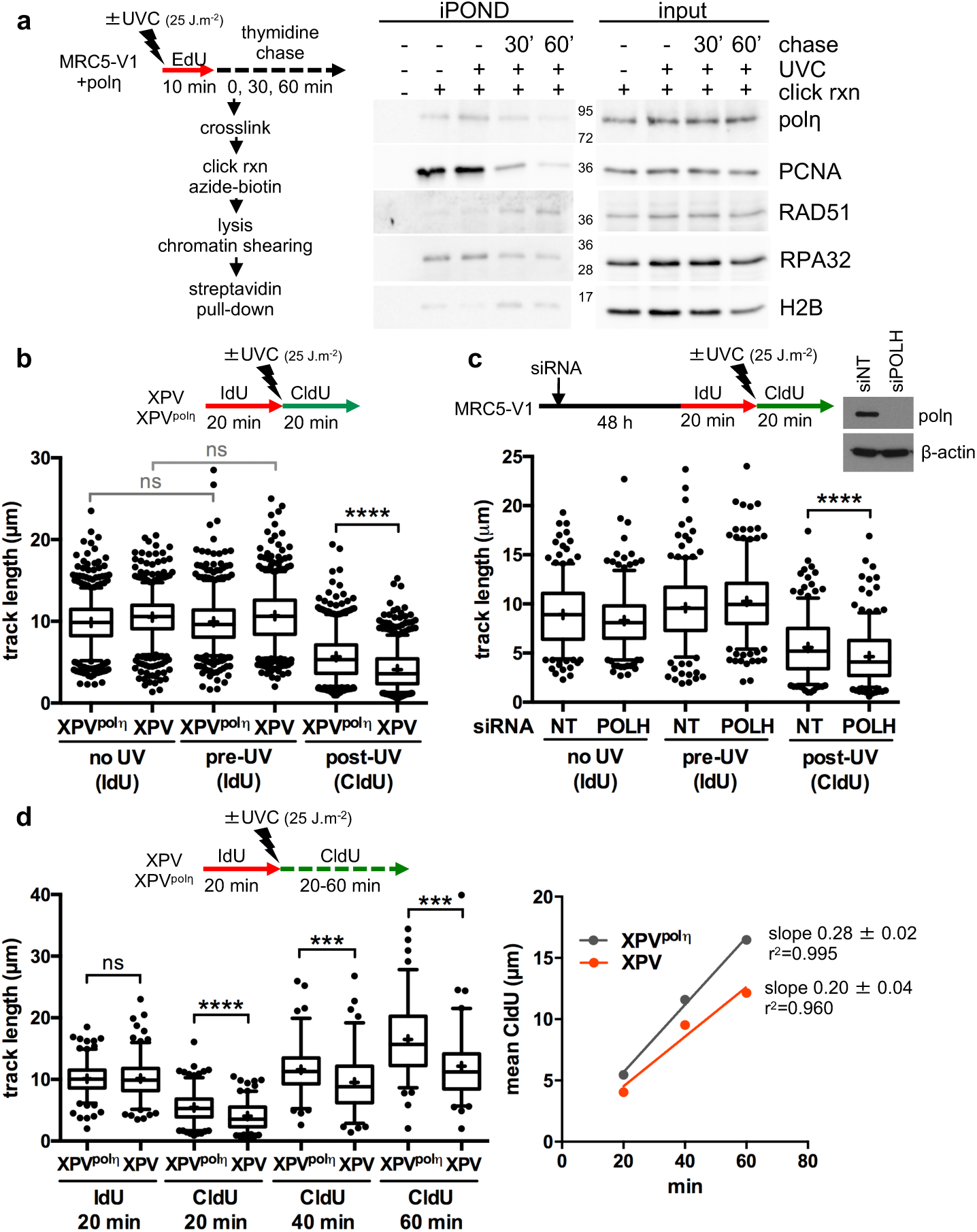
Polη promotes replication fork progression after UV. (**a**) MRC5-V1 cells stably expressing pol*η* were irradiated or not with UVC (25 J.m^-2^) and pulse-labelled with 10 μM EdU for 10 min. Cells were cross-linked immediately after the pulse or after a 30- or 60-min thymidine chase. Biotin was conjugated to EdU by click chemistry (click rxn) prior to cell lysis and chromatin shearing. Labelled DNA was retrieved on streptavidin beads and input and bound proteins (iPOND) were analysed by western blot using the indicated antibodies. (**b**) XPV cells and their corrected counterpart (XPV^polη^) were irradiated or not at 25 J.m^-2^ between 20-min pulses of IdU and CldU. DNA was spread on glass slides and IdU, CldU and total DNA were detected by immunofluorescence. Replicated tracks were measured on unbroken bicolor signals (pool of 3 independent experiments, n>750, box-plot with interquartile range and 5-95 percentile whiskers, +: mean of the distribution, dots: outliers). (**c**) MRC5-V1 cells were depleted for pol*η* using siRNAs and treated like in b (pool of 2 independent experiments, n=300, siNT: non specific siRNA, siPOLH: siRNA against *POLH* mRNAs). siRNA efficiency was confirmed by western blot. (**d**) XPV and XPV^polη^ cells were treated as in b but with a CldU pulse of 20, 40 or 60 min. Left panel shows the distribution of IdU and CldU tracks (n>90). Right panel shows the linear regression of the mean CldU calculated from this distribution. Slopes and r^2^ coefficients are indicated. Data were analysed for statistical significance with the Mann-Whitney test (ns: not significant, ***: P<0.001, ****: P<0.0001).

To determine if pol*η* deficiency impacts RF progression in presence of UV lesions, we performed DNA fiber assays in XPV cells and their counterpart stably complemented with wild-type pol*η* (herein referred as XPV^polη^ cells). Cells were irradiated between two pulses of iododeoxyuridine (IdU) and chlorodeoxyuridine (CldU) and unbroken bicolour signals were measured to compare the distances covered before and after UV exposure at the single-fork level. Replication tracks were shorter after UV and this defect was further aggravated by pol*η* deficiency (Figure 1b). Similar results were found by inverting the two thymine analogues (Supplementary Figure 1a) or after depletion of endogenous pol*η* in SV40-immortalized human MRC5-V1 fibroblasts (Figure 1c). This shows that pol*η* promotes RF progression in presence of UV lesions, in agreement with previous findings^24–26^. Moreover, we showed that this required pol*η* catalytic activity, as well as its PCNA- and ubiquitin-binding motifs, two regulatory domains that cooperate for pol*η* accumulation in replication foci^36^ (Supplementary Figure 1b). Increasing the duration of the post-UV pulse led to longer tracks in both cell lines (Figure 1d). Linear regression analysis showed that RFs progress at an apparent constant average speed in the first hour following irradiation, this speed being lower in pol*η*-deficient cells. Moreover, forks presenting a severe slow down (defined as post-UV length lower than 30% of the pre-UV length) were more abundant in XPV cells than in XPV^polη^ cells 20 min after UV, but they did not persist at later time points (Supplementary Figure 1c). Hence, RFs are not permanently stalled in irradiated XPV cells where alternative restart mechanisms can take place, although with a decrease efficiency and/or with a delay.

In pol*η*-deficient cells, residual error-prone TLS was shown to involve two pathways depending on pol*ζ* and pol*κ*, respectively^9–11^. However, depletion of pol*κ* and/or of REV3L, the catalytic subunit of pol*ζ*, did not further aggravate the RF progression defect observed in XPV cells (Supplementary Figure 1d), suggesting that, although required for XPV cell survival and for UV-induced mutagenesis, these alternative polymerases act on a different substrate than stalled RFs.

### Polη limits ssDNA accumulation at and behind replication forks after UVC

To determine which mechanisms sustain RF progression in absence of pol*η*, we analysed the architecture of RFs by electron microscopy at high resolution. Cells were irradiated or not with UVC and subjected to UVA/psoralen *in vivo* crosslinking to stabilize replication intermediates (RIs) by avoiding branch migration. Purified DNA was enriched in RI-containing material as already described^5, 39^. To characterize the various DNA intermediates structures, we developed a procedure which combines BAC methods and positive staining to optimize the recruitment and the spreading of DNA onto the carbon film and to provide an optimal contrast and resolution using darkfield imaging mode^40^.

As expected, in mock-irradiated samples, the majority of RFs displayed no or limited ssDNA stretches at the elongation point (Figure 2a), presumably on the lagging strand, as previously shown^5, 41^. Thirty min after UVC irradiation (25 J.m^-2^), two kinds of consequences were observed. First, some RFs displayed an extended ssDNA stretch at one side of the elongation point (Figures 2b,d,e,g), corresponding to the blockage of the leading strand synthesis and its subsequent uncoupling from the helicase activity, as previously observed by TEM on plasmid models^41^ and later confirmed in yeast and human *in vitro* and *in vivo* studies^5, 18, 33, 42^. Secondly, some RFs showed single-stranded regions behind the elongation point, on one or both daughter strands (Figures 2c-i). Some RIs displayed successive gaps on the same strand (Figure 2h), gaps in opposite strands (Figure 2f) or both uncoupling and post-replicative gap(s) (Figures 2d,e,g), presumably because UV lesions are formed at a higher frequency in peculiar genomic regions^43^. Very long internal gaps were rarely observed (Figures 2i). Unlike what was observed by others at a later time point^33^, we did not find evidence of fork reversal or recombination-like processes early after UVC irradiation in our conditions.

**Figure 2.**
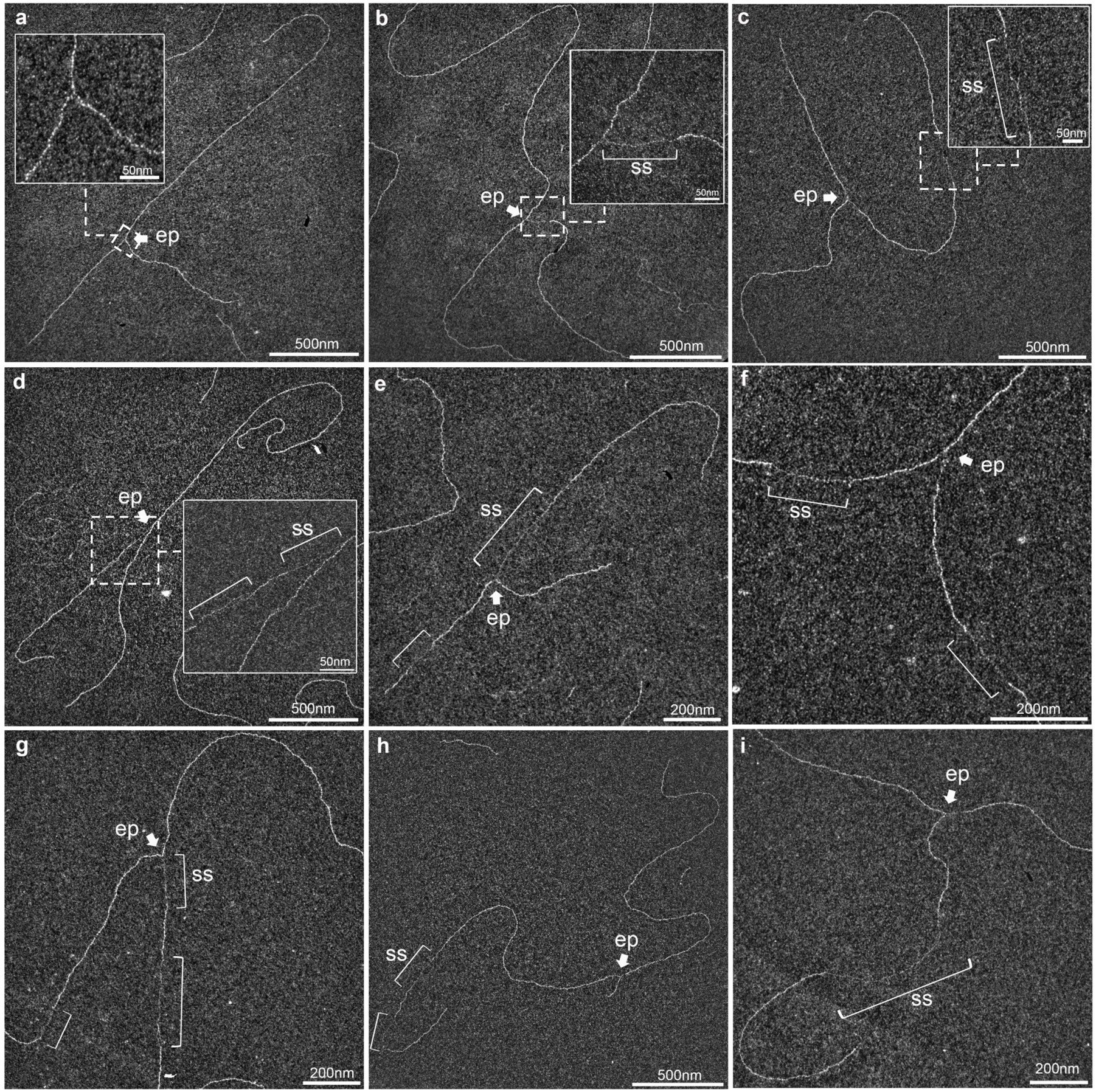
UVC irradiation leads to long ssDNA stretches at the elongation point and to post-replicative gaps as analysed by TEM in annular dark field mode. (**a**) Usual replication fork arising from the hydrolysis of a replication bubble by enzymatic restriction. The fork does not display ssDNA regions and is symmetric as regards to the identical daughter strand lengths. (**b-i**) Replication intermediates observed 30 min after exposure to UVC (25 J.m^-2^). Inlays represent enlargements of the regions framed with dotted lines (scale bar = 50 nm). Panels b, d, e and g show uncoupled replication forks with ssDNA starting from the elongation point because of DNA synthesis block on the leading strand. Panel c shows a fork with an internal ssDNA gap on one daughter strand. The gap can appear on the leading strand (d), the lagging strand (e) or both (f, g). Gaps can accumulate successively on the same daughter strand (h). Forks displaying ssDNA at and behind the elongation point can also be observed (d, e, g). Unusually long gaps are rarely observed (i, internal gap length = 545 nm). ep: elongation point; ss: ssDNA.

To allow quantitative comparison of pol*η*-proficient and pol*η*-deficient cells, ssDNA regions were measured on more than 50 RFs in two independent experiments. UVC led to a significant increase of the ssDNA median length both at and behind RFs in XPV cells compared to XPV^polη^ cells (Figures 3a,b). Given that short ssDNA regions were also observed in untreated conditions, we applied a threshold to consider only long ssDNA resulting from UVC exposure. We chose the 95^th^ percentiles of the distributions of untreated XPV^polη^ cells. Interestingly, these thresholds were similar at and behind the forks (84 nm and 85 nm, respectively), and correspond to a distance of approximately 165 nucleotides, close to the size of an Okazaki fragment^44^. Consequently, RFs bearing ssDNA at the elongation point above the threshold likely correspond to uncoupled forks. After UVC irradiation, XPV cells showed a 4-fold increase of ssDNA at RF compared to XPV^polη^ cells (Figure 3c), in agreement with the stronger slow down observed in DNA fiber assays. RFs with UV-induced post-replicative gaps were 2.4 times more frequent in pol*η*-deficient cells (Figure 3d), which also showed more gaps per RF (Supplementary Figure 2a). In irradiated XPV cells, 47.2% of forks were uncoupled and 57.7% were gapped, with structures cumulating both particularities representing 22.8% (vs 2.6% in XPV^polη^ cells) (Supplementary Figure 2b), suggesting that most of the forks have encountered at least one lesion in our experimental conditions.

**Figure 3.**
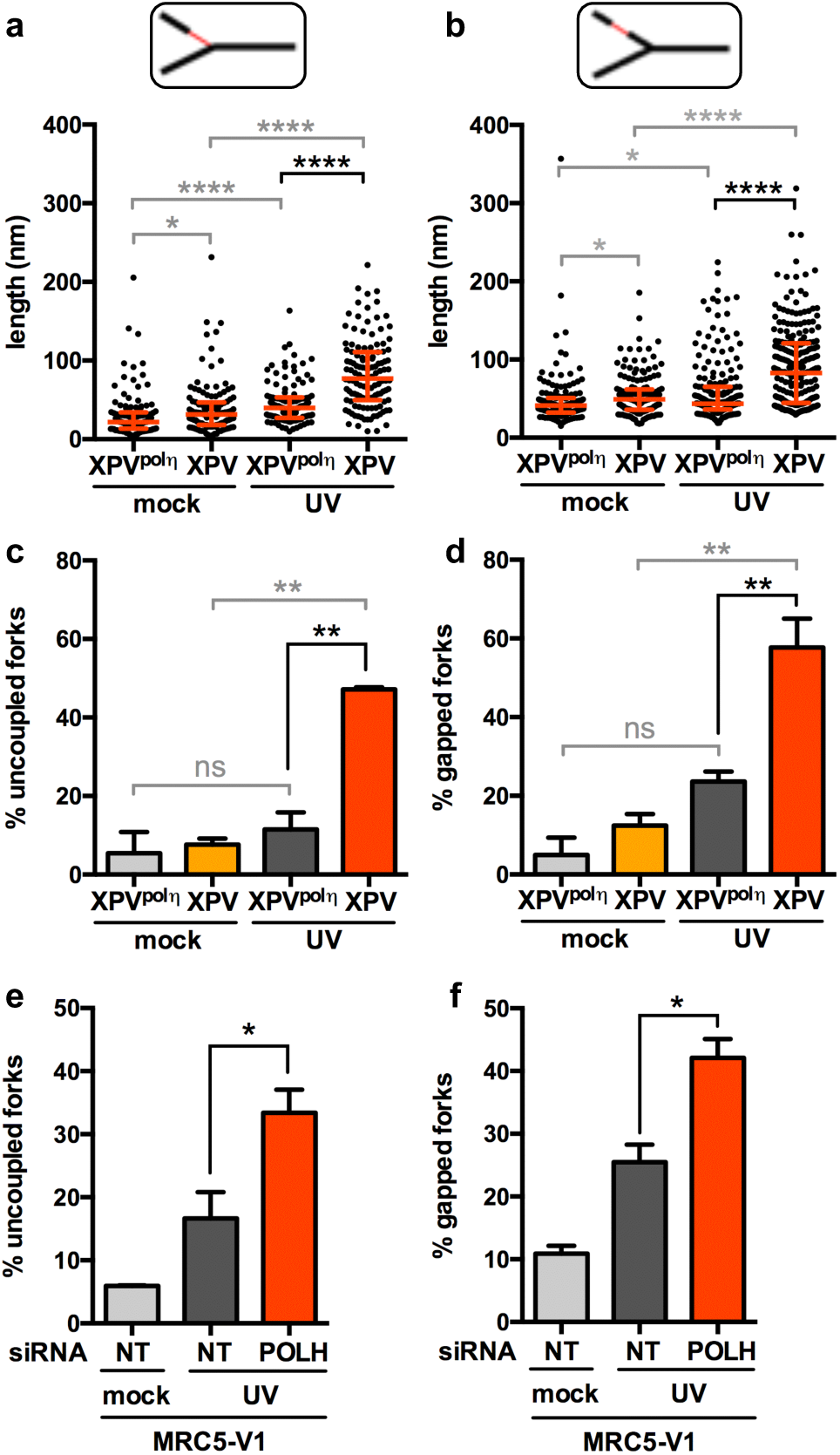
Polη prevents accumulation of ssDNA at and behind the elongation point after UV. TEM analysis was performed in XPV and XPV^polη^ cells 30 min after UVC irradiation at 25 J.m^-2^ in two independent experiments. For each condition, at least 50 forks were analysed per experiment. (**a**) Distribution of the lengths of the ssDNA regions located at RFs (pool from the 2 experiments, n>100). (**b**) Distribution of the lengths of the ssDNA regions located behind RFs. Data are shown as dot plots with medians and interquartile ranges. Statistical significance was tested with the Kruskal-Wallis test (*: P<0.05, ****: P<0.0001). The 95^th^ percentiles of the distributions of untreated XPV^polη^ cells (84.23 nm and 85.26 nm at and behind RFs, respectively) were used as a threshold to identify UV-induced long ssDNA tracks at (uncoupling) and behind (gap) the elongation point. (**c**) Proportion of uncoupled forks. (**d**) Proportion of gapped forks. Values are the mean ± SD of the two independent experiments. Statistical significance was tested by ordinary Anova (ns: not significant *: P<0.05, **: P<0.01). The same procedure was performed in MRC5-V1 cells depleted or not for pol*η*. (**e**) Proportion of uncoupled forks and (**f**) proportion of gapped forks were determined as described above.

Interestingly, ssDNA at the elongation point rarely exceed 450 nucleotides, even in pol*η*-deficient cells, suggesting that uncoupling between helicase and stalled leading strand synthesis is spatially limited. The distributions of UV-specific post-replicative gap lengths are similar in XPV and XPV^polη^ cells (Supplementary Figure 2c), supporting a common mechanism for gap formation in presence or absence of pol*η*. Moreover, this suggests that the gaps formed in XPV^polη^ cells are unlikely to be filled in in a pol*η*-dependent manner in the first 30 min following UV exposure, since gap lengths should then have been statistically reduced compared to XPV cells. A similar conclusion was made in yeast^5^. Assuming that long ssDNA at the elongation point signals an uncoupled leading strand, we could discriminate strands on 56 RIs in the UV-irradiated XPV samples, 27 of which did not present UV-specific gaps. On the other 29, we counted 17 and 29 gaps on the leading and the lagging strands, respectively. Hence, although lagging strand gaps were more likely to occur and were found in 39.3% of the molecules included in this analysis, gap formation on the leading strand was not a rare event and was observed in 23.2% of the RIs. Dynamic accumulation of ssDNA at and behind RFs in XPV cells was associated with an enhanced recruitment of the ssDNA-binding complex RPA on chromatin (Supplementary Figure 2d), as previously described^24^. Interestingly, while the RPA signal remained elevated up to 24 h in XPV cells after 25 J.m^-2^, in XPV^polη^ cells it peaked at 2-5 h, then slowly decreased over 20 h. We assumed that this decrease corresponds to repair/gap-filling events, which will be further characterized.

Importantly, these findings were recapitulated in MRC5-V1 fibroblasts upon depletion of pol*η* by siRNAs (Figures 3e,f and Supplementary Figures 2e-g), confirming that our observations are entirely dependent on pol*η* loss.

Altogether, our data show that pol*η*-dependent TLS at RFs is a crucial mechanism to sustain a continuous DNA synthesis, preventing accumulation of uncoupled forks and post-replicative ssDNA gaps. Altered forks were also observed in pol*η*-proficient cells to a lesser extent, suggesting that a subset of UV lesions cannot be bypassed at RFs even when pol*η* is functional. These lesions are presumably the 6-4PPs, which represent 20-25% of the UV-induced DNA lesions and cannot be bypassed by pol*η* alone^25^. Moreover, our data support a higher occurrence of repriming events when pol*η* is not functional.

### PrimPol sustains RF progression in XPV cells but is not the only actor of repriming downstream UV lesions

Actors allowing the repriming of DNA replication downstream DNA damage remain elusive in eukaryotes. Recent studies revealed that the primase activity of PrimPol, an archaic primase and TLS polymerase, mediates fork progression after UV and hydroxyurea (HU) in vertebrates^20–22^. However, to date, no structural data fully support this model. Moreover, PrimPol invalidation does not lead to dramatic sensitization to UV light, even when pol*η* is not functional^45^ (and data not shown).

First, we showed that PrimPol, but not the catalytic polymerase sub-unit of polα POLA1, was recruited on chromatin as early as 30 min after UVC and to a higher extent in XPV cells compared to XPV^polη^ cells (Figure 4a, Supplementary Figure 3a), suggesting an increased usage of PrimPol in absence of pol*η*. DNA fiber assays showed that PrimPol depletion impedes RF progression after UV to a higher extent in XPV cells (Figure 4b) than in their complemented counterpart (Supplementary Figure 3b), in agreement with previous findings^45^. However, even in XPV cells, the impact of PrimPol deficiency was moderate in the first 20 min following UVC irradiation and became prominent in the 20-40 min time window further analysed, with a significant 2-fold decrease in the mean fork rate and the persistence of forks presenting a severe slow down (Figure 4b). Intriguingly, PrimPol depletion did not impact on RPA recruitment on chromatin early after UV irradiation whether pol*η* is present or not (Figure 4c).

**Figure 4.**
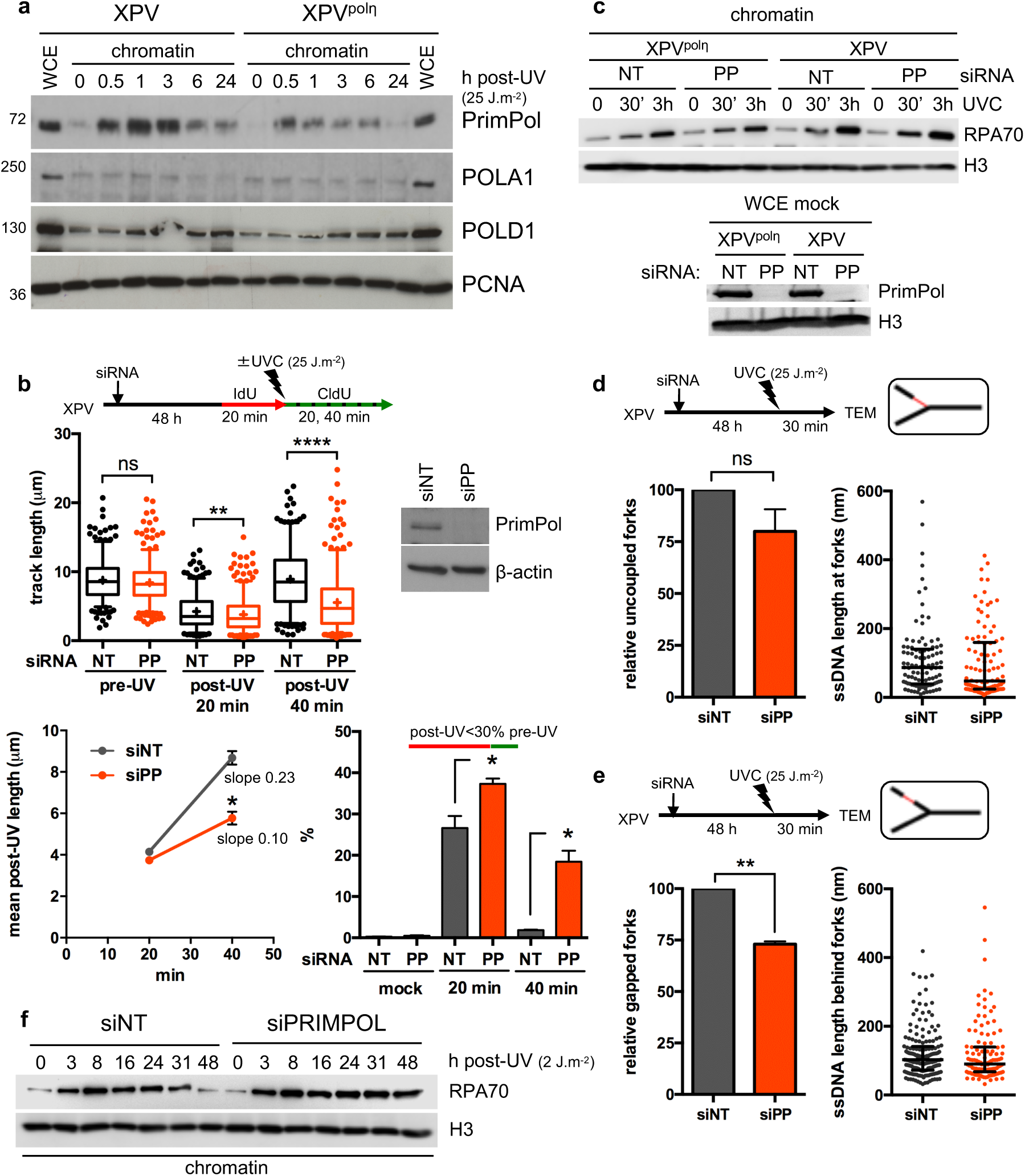
PrimPol depletion further reduces replication track length and impacts on RI architecture in XPV cells. (**a**) Cellular fractionation was performed at the indicated time points following UVC irradiation at 25 J.m^-2^ in XPV and XPV^polη^ cells. Whole cell extracts (WCE) of unirradiated cells and chromatin-bound proteins were analysed by western blot using the indicated antibodies. (**b**) XPV cells were depleted for PrimPol (PP) and DNA fiber assay was performed as described in Figure 1d. Upper panel show the distributions of replication track length (n>300, Mann-Whitney test, ns: not significant; **: P<0.01, ****: P<0.0001). siPRIMPOL efficiency was confirmed by western blot. An independent experiment was performed with CldU and IdU as pre- and post-UVC labelling, respectively, and gave the same results. The middle and lower panels show the average post-UV length and the proportion of replication signals with post-UV length shorter than 30% of the corresponding pre-UV length for these two experiments, respectively (mean ± SD, t-test, *: P<0.05). (**c**) Upper panel: kinetics of recruitment of RPA70 on chromatin after UVC (25 J.m^-2^) in XPV and XPV^polη^ cells upon PrimPol depletion. Lower panel: WCEs were used to confirm the efficiency of PrimPol depletion. (**d**) Relative uncoupled forks and distribution of ssDNA lengths at the elongation point in irradiated XPV cells following PrimPol depletion. (**e**) Relative gapped forks and distribution of gap lengths in irradiated XPV cells following PrimPol depletion. At least 50 forks were characterized in two independent experiments. Histograms are the mean ± SD of the two experiments (t-test, ns: not significant, **: P<0.01). ssDNA length distributions are pools of the two experiments (dot plot with median and interquartile range). (**f**) Kinetics of chromatin-bound RPA70 fraction in PrimPol depleted XPV cells after irradiation at a low UVC dose (2 J.m^-2^).

If PrimPol indeed contributes to RF progression through its repriming activity, its depletion should lead to increased fork uncoupling and decreased post-replicative gaps, consequences expected to be enhanced in XPV cells. TEM analysis did not reveal any significant impact of PrimPol depletion on RI architecture in pol*η*-expressing cells 30 min after UV (Supplementary Figure 3c). In XPV cells, the median ssDNA length at the elongation point unexpectedly decreased when PrimPol was depleted (Figure 4d). Although this difference was not statistically significant, it was observed in two independent experiments (data not shown) and was not observed in pol*η*-expressing cells (Supplementary Figure 3c). One simple explanation could be that decreased RF rate observed upon PrimPol depletion reduces the chance to encounter a UV lesion, mitigating RF uncoupling. Alternatively, remodelling of some RIs might occur in absence of both pol*η* and PrimPol. In support with this latter hypothesis, we observed a transient decreased of PCNA and other replisome components on chromatin in PrimPol-depleted XPV cells (Supplementary Figure 3d).

Although the overall number of internal gaps is lower in absence of PrimPol (64 gaps vs 111 gaps on the 102 RFs analysed in siPRIMPOL and siNT cells, respectively), numerous post-replicative gaps still formed in PrimPol-depleted XPV cells, with similar gap length distributions than in mock-depleted cells (Figure 4e and Supplementary Figure 3e). PrimPol depletion led to a 25% decrease in the proportion of gapped RFs (Figure 4e). Interestingly, this reduction seems specifically related to the RIs bearing both uncoupling and internal gaps (Supplementary Figure 3f), suggesting that PrimPol primase activity may deal with regions bearing a high density of damage. On this kind of RIs, the extended ssDNA at the elongation point allows identifying the leading strand. We observed that PrimPol depletion preferentially impaired gap formation on the leading strand (Supplementary Figure 3g), which suggests that PrimPol can indeed contribute to reprime some stalled leading strands in XPV cells. However, although siRNAs against PrimPol were highly efficient (Figure 4c), this impairment is only partial, indicating that PrimPol-independent mechanisms also occur.

Hence, our data suggest that PrimPol primase activity is required in specific conditions, i.e high density of UV-induced lesions and unavailability of pol*η*-dependent bypass at the RF. The global effect of PrimPol depletion on RF progression observed in DNA spreading assay may therefore reflect the involvement of its TLS polymerase or lesion skipping activities. Moreover, we cannot rule out the possibility that impaired mitochondrial DNA replication induced by PrimPol deficiency indirectly contributes to the impaired genome replication in presence of DNA damage^21, 45, 46^.

To determine if PrimPol depletion could hamper post-replicative gap filling, we treated PrimPol-depleted XPV cells with a sub-lethal UV dose (2 J.m^-2^), to be able to monitor RPA amounts on chromatin at later times after irradiation (Figure 4f). We have previously shown that this dose leads to a strong reduction of DNA synthesis in XPV cells during the first 24 hours and that normal levels are restored after 48 h^24^. In these conditions, the chromatin-bound RPA amount peaked around 8 h and went down to the unirradiated levels between 30 and 48 h after UV exposure in cells transfected with a non-targeting siRNA (Figure 4f). In contrast, it remained elevated in PrimPol-depleted XPV cells. This suggests that PrimPol could also contribute to gap repair.

### Different Rad51 activities act at RFs and in post-replicative gap filling

In addition to its role in DSB repair, HR was proposed to restart stalled RFs and to contribute to fill in post-replicative gaps both in yeast and mammalian cells^5, 32, 47, 48^. Moreover, HR was proposed to act earlier and to a greater extent in irradiated XPV cells than in pol*η*-proficient cells to compensate for pol*η*deficiency^13^. However, it is not clear if this earlier commitment drives RF progression or results from the accumulation of post-replicative gaps in absence of pol*η*.

Given that we observed an accumulation of RAD51 behind RFs in pol*η*-expressing cells (Figure 1a), we first compared the localisation of chromatin-bound RAD51 with respect to RFs in XPV^polη^ and XPV cells. iPOND confirmed that UV irradiation specifically enhanced RAD51 behind RFs in XPV^polη^ cells, in wild-type human MRC5-V1 fibroblasts or in wild-type mouse embryonic fibroblasts (Figure 5a and Supplementary Figures 4a,b), indicating a post-replicative commitment of RAD51 when pol*η* is functional, presumably in gap repair. The amount of RAD51 in damaged post-replicative chromatin was further enhanced in XPV cells that also displayed more residual PCNA behind RFs and a defect in chromatin restoration, as evidenced by an impaired incorporation of histone H3 (Figure 5a), in agreement with the accumulation of post-replicative gaps observed by TEM. Interestingly, XPV cells also displayed a premature recruitment of RAD51 in the vicinity of RFs (Figure 5a). This was confirmed by proximity ligation assay (PLA) between RAD51 and nascent DNA (Supplementary Figure 4c). Importantly, a similar RAD51 dynamics was observed after irradiation of XPV cells at a lower UVC dose (Supplementary Figure 4d).

**Figure 5.**
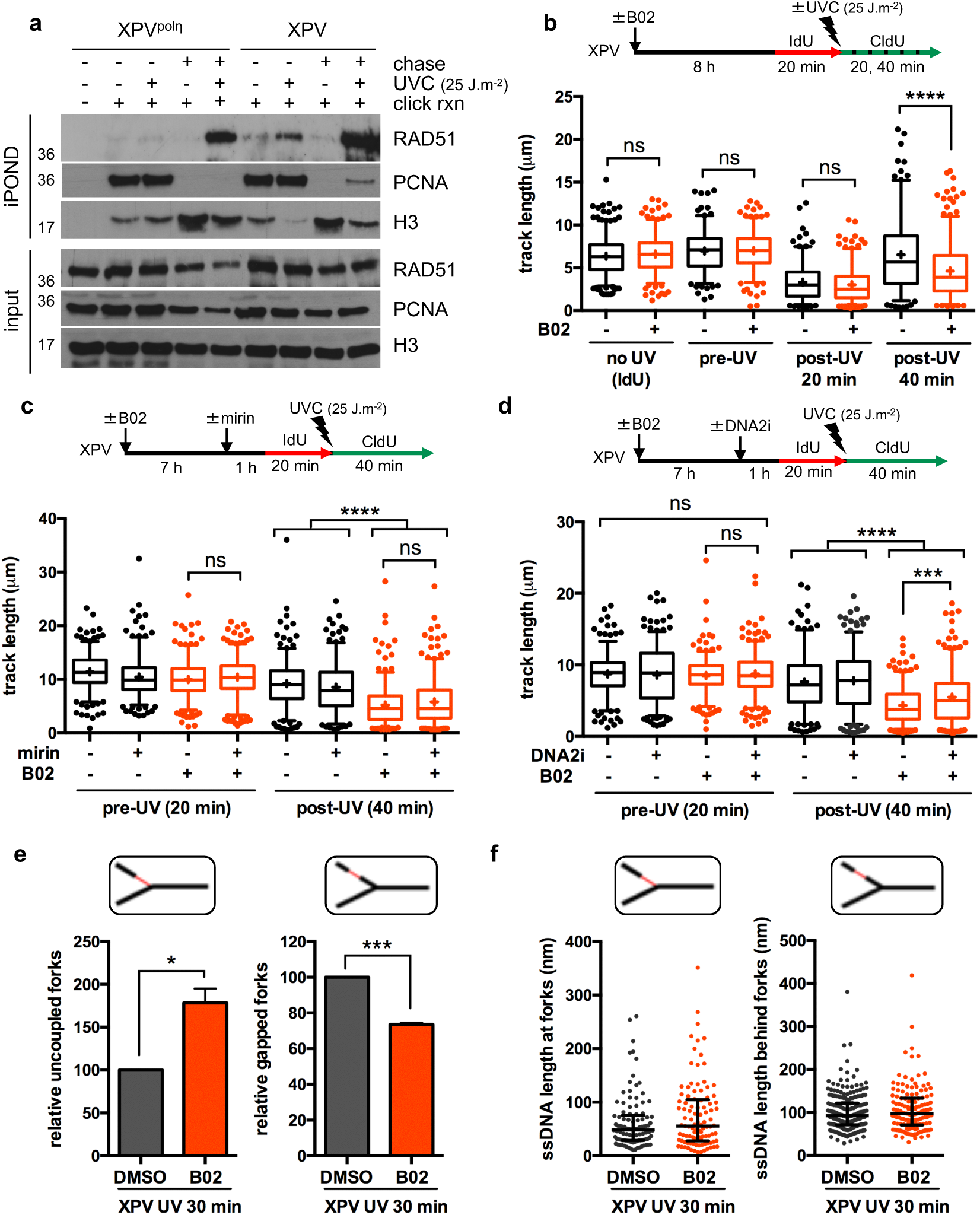
RAD51 solves RF uncoupling in UV-irradiated XPV cells. (**a**) iPOND was performed in XPV and XPV^polη^ cells as described in Figure 1a with a 1-hour thymidine chase when indicated. Immunoblotting was done with the indicated antibodies. (**b**) DNA fiber assay was performed in XPV cells pre-treated or not with 20 μM B02 for 8 h. The distributions of replication track lengths are shown (n>200, Mann-Whitney test, ns: not significant; ****: P<0.0001). (**c**) XPV cells were pre-treated with 20 μM B02 for 8 h and 50 μM MRE11 inhibitor MIRIN for 1 h (n>250, Mann-Whitney test, ns: not significant; ****: P<0.0001). (**d**) XPV cells were pre-treated with 20 μM B02 for 8 h and 25 μM DNA2 inhibitor C5 (DNA2i) for 1 h (n>250, Mann-Whitney test, ns: not significant; ***: P<0.001; ****: P<0.0001). (**e**) The relative numbers of uncoupled and gapped forks were determined by TEM 30 min after UVC exposure (25 J.m^-2^) in XPV cells pre-treated or not with 20 μM B02 for 8 h (mean ± SD of two independent experiments, at least 50 RIs were analysed per condition in each experiment, t-test, *: P<0.05, ***: P<0.001). (**f**) The distributions of the lengths of ssDNA measured at and behind RFs in the TEM experiments (pool) are shown as dot plots with median and interquartile range.

To further dissect the functions of RAD51 at and behind RFs, we used the RAD51 inhibitor B02, which inhibits the binding of RAD51 on DNA and, subsequently, its strand-exchange activity^49^. In unirradiated XPV cells, we observed a decrease of the chromatin-bound RAD51 6 h post-B02 treatment (Supplementary Figure 4e), in agreement with previous studies reporting a delay of B02 activity^49, 50^. To determine the consequences of RAD51 inhibition early after UV, we pre-treated XPV cells with B02 for 7-8 h prior to UV exposure and confirmed by immunofluorescence that this impaired UV-induced recruitment of RAD51 on chromatin in S phase cells (Supplementary Figure 4f). In these experimental conditions, B02 treatment led to shorter post-UV replication tracks, with no impact on pre-UV tracks or in unirradiated cells (Figure 5b). While the restoration of post-UV track lengths was not observed after incubation with the MRE11 3’-5’ exonuclease activity inhibitor Mirin (Figure 5c), it was partially achieved upon inhibition of the 5’-3’ DNA2 nuclease (Figure 5d), suggesting that shorter post-UV tracks were not the consequence of excessive neo-synthesised DNA degradation. Analysis of RF architecture by TEM showed that B02 treatment increased the proportion of uncoupled RFs while it simultaneously reduced the occurrence of post-replicative gaps (Figure 5e). ssDNA length distributions remained similar to that of DMSO-treated cells, confirming that RAD51 inhibition did not induce extensive resection at or behind RFs early after UV (Figure 5f). Altogether, these data raise the possibility that RAD51 acts at RFs to promote repriming in absence of pol*η*on-the-fly bypass. This function may partially depend on protection of the reprimed leading strand towards DNA2 activity (Figure 5d).

To confirm these findings, we knocked down RAD51 by RNA interference in XPV cells. Of note, since it drastically impaired cell growth and led to tetraploidization in our cellular model (data not shown), we lowered the amount of siRNAs and performed our experiments in the first 36 hours following transfection. Similarly to B02 treatment, RAD51 depletion led to shorter post-UV replication tracks, a phenotype partially dependent on DNA2 but not MRE11 activity (Supplementary Figures 5a,b). This was associated with a progressive attrition of pre-UV replication tracks, as already described^26^, which was counteracted by DNA2 or MRE11 inhibition. Using TEM, we observed a strong reduction in the proportion of RIs bearing post-replicative gaps in RAD51-depleted XPV cells (Supplementary Figure 5c), supporting impaired repriming. Moreover, a global decrease in ssDNA stretch lengths at RFs was observed (Supplementary Figure 5d), suggesting that acute loss of RAD51 at damaged RFs may result in fork backtracking, whereas uncoupled structures are maintained upon partial inhibition of RAD51 activity by B02. Noteworthy, the impact of RAD51 depletion is less severe in pol*η*-proficient cells, specifically affecting RIs with both gaps and uncoupling (Supplementary Figures 5e,f). Once again, this supports the key role of pol*η* at UV-stalled replication forks.

Interestingly, TEM analysis performed 2 h after UV exposure (25 J.m^-2^) revealed unusual multi-branched (or H-shaped) structures, usually bearing ssDNA stretches, in both XPV and XPV^polη^ cells (Figure 6a). In XPV cells, these structures represent slightly more than 10% of the observed intermediates (5 and 7 H-shaped structures for 50 forks counted in two independent experiments) while none were observed in unirradiated XPV samples. We assume that these multi-branched structures most likely corresponded to HR specific strand exchange intermediates since TEM analysis of joint molecules formed in the *in vitro* RAD51-dependent strand-exchange assay revealed the formation of exactly the same type of structures (Figure 6b). Interestingly, the multi-branched intermediates resulted from the existence of an ssDNA segment (97.7 ± 19 nm long, about 189 nucleotides) at the point of strand transfer between DNA molecules. We then assessed the post-replicative role of RAD51 by adding B02 directly after irradiation at a low UVC dose (2 J.m^-2^). This UV dose is too low to induce an increase of chromatin-bound RPA in pol*η*-proficient cells. Whereas B02 treatment did not affect RPA accumulation on chromatin 3 h and 8 h after irradiation in XPV cells, it impaired RPA disappearance at later time points in a dose-dependent manner (Figure 6c and Supplementary Figure 6a). To a lower extent, we observed a similar effect in XPV^polη^ cells at a higher UVC dose (Supplementary Figure 6b). This suggests that RAD51 is required for efficient post-replicative gap repair. We therefore hypothesised that the post-replicative recruitment of RAD51 and the multi-branched structures observed *in vivo* correlate and reflect post-replicative recombination events occurring to fill in the gaps accumulated behind the RFs.

**Figure 6.**
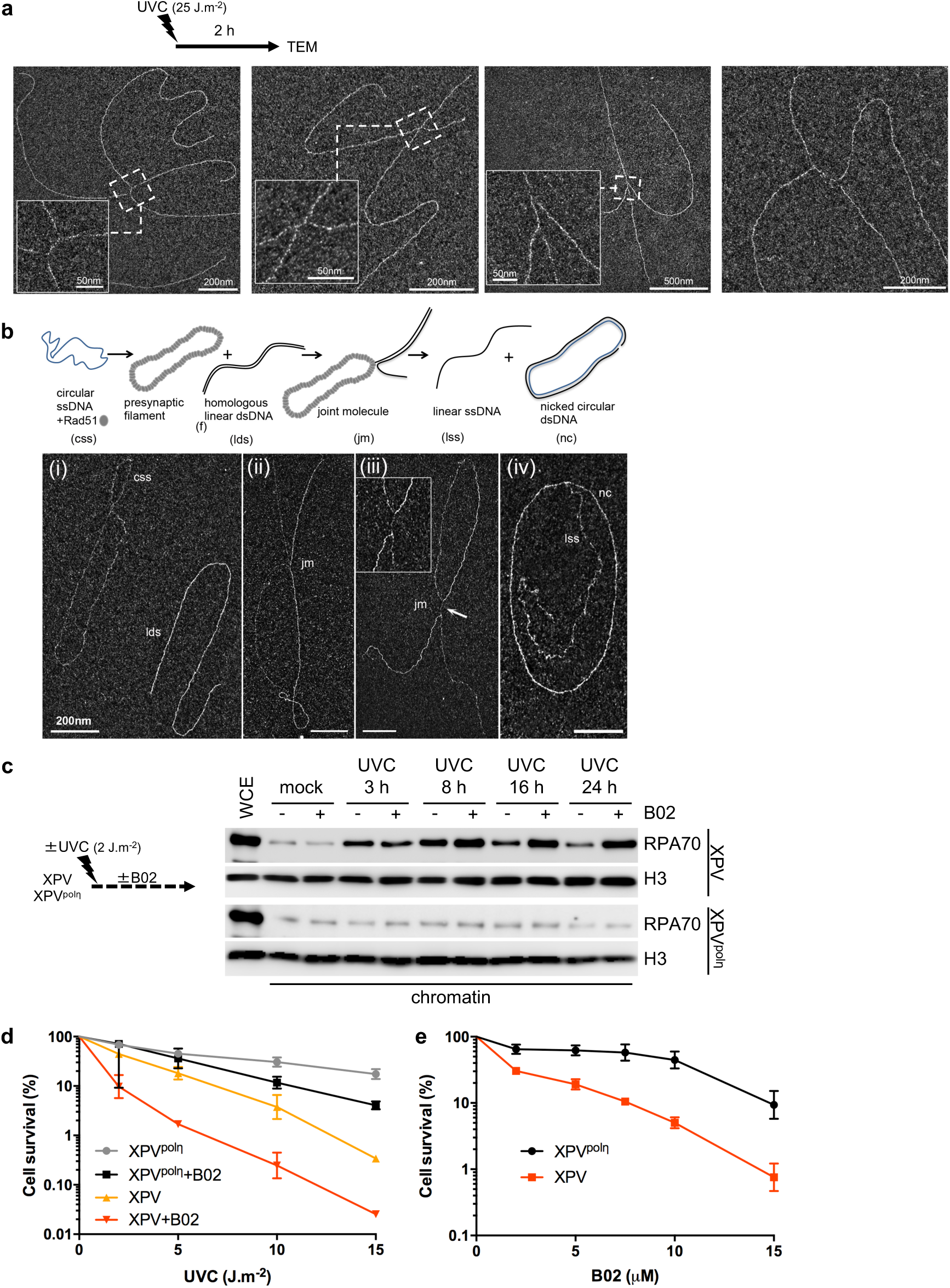
RAD51 is required for post-replicative gap filling and cell survival. (**a**) Examples of *in vivo* five-branched structures observed by TEM 2 h after UVC (25 J.m^-2^). Inlays represent enlargements of the regions framed in dotted lines (scale bar = 50 nm). (**b**) Examples of joint molecules formed in the *in vitro* RAD51-dependent strand-exchange assay observed by TEM. The circular ssDNA Phix174 virion was incubated with RAD51, followed by RPA addition, then linearized Phix174 RFI homologous plasmid was added to the reaction. RAD51 polymerizes onto the ssDNA substrate to form the presynaptic filament and then catalyses the strand invasion and strand transfer with the linear homologous dsDNA. Final products are linear ssDNA (lss) and nicked dsDNA plasmid (nc). For TEM observations, reactions were subjected to UVA/trioxalen *in vitro* crosslinking, deproteinization, and spread using the BAC hyperphase method. (i) circular ssDNA Phix174 virion (css) and linearized homolog plasmid (lds). (ii), (iii) Representative views of joint molecules. Formation of five-branched (H-shaped) structures is shown in (iii). (iv) Final products of the reaction. Scale bar = 200 nm. (**c**) XPV and XPV^polη^ cells were irradiated with UVC at 2 J.m^-2^. 10 μM B02 was added immediately after irradiation. RPA70 recruitment to chromatin was assessed at different times by cellular fractionation. (**d**) XPV and XPV^polη^ cells were irradiated with increasing UVC doses. When indicated, 10 μM B02 was added post-irradiation. Cell survival was assessed 72 h after treatment (mean ± SD of three independent experiments. For each condition, 100% represents the number of cells in the non-irradiated conditions). (**e**) Cell survival following 2 J.m^-2^ UVC upon increasing concentrations of B02 (mean ± SD of four independent experiments. For each cell line, 100% represents the number of cells in the DMSO condition).

The effect of RAD51 inhibition on cell proliferation was similar in unirradiated XPV and XPV^polη^ cells (data not shown). However, although B02 treatment sensitized both XPV and XPV^polη^ cells to UVC, its impact was stronger in pol*η*-deficient cells (Figures 6d,e). In particular, at the sub-lethal dose of 2 J.m^-2^, XPV cells were sensitized by lower concentrations of B02 than XPV^polη^ cells (Figure 6e). Importantly, these findings were fully recapitulated in pol*η*-depleted MRC5-V1 cells (Supplementary Figures 6c,d).

Altogether, our data suggest that RAD51 acts in two different mechanisms during replication of UV-damaged DNA. First, it favours RF progression through repriming when pol*η*-mediated TLS cannot ensure on-the-fly bypass. As we did not observe recombination-like structures early after UV, we hypothesise that this function does not involve RAD51 strand-invasion activity but rather protective function of RAD51 towards DNA2 activity. Secondly, it accumulates behind RFs to allow recombination-based post-replicative gap-filling independently of the pol*η* status. As XPV cells more strongly rely on repriming and gap filling for completion of DNA replication, this consequently results in an important dependence of XPV cells towards these two RAD51 activities.

## Discussion

The present time-course characterization of RIs architecture by TEM, combined with biochemical approaches, provides new insights for the understanding of the organisation of DDT after exposure of human cells to UV irradiation. From our data, we propose that pol*η*-dependent TLS is the primary and main fork restart mechanism and that its efficiency configures the DDT response. When this pathway is not possible, either at UV lesions that cannot be bypassed by pol*η* alone or because pol*η* is not functional, DDT is diverted to a post-replicative mechanism through repriming. Interestingly, we also revealed that RAD51 participates in both the repriming and the fill in of the post-replicative gaps generated by repriming and that only this second function involves strand exchange transactions.

### Impact of UV damage and polη deficiency on the architecture of replication forks

Our TEM analysis showed that the early response of RFs to UV damage is the generation of abnormally long ssDNA regions at and behind the elongation point, the amount of which is related to the capacity of pol*η* to bypass lesions in the vicinity of RFs. Thanks to the unique structure of its catalytic site, pol*η* ensures efficient and accurate bypass of TT-CPDs^8^. Furthermore, we have previously shown that pol*η* travels with RFs during unperturbed S phase^38^ and here we show that it further rapidly accumulates at RFs after UV. Hence, replisomes of WT cells possess the enzyme allowing the on-the-fly bypass of the most abundant UV-induced lesion and, therefore, a limited impact of UV damage on RF architecture, whereas pol*η* deficiency leads to RF uncoupling and accumulation of ssDNA gaps in both lagging and leading strands.

In yeast reconstituted replisome, the stalling of the leading strand synthesis at a single lesion leads to the uncoupling of the replicative helicase and the slowdown of DNA unwinding^18, 51^. Our data support this model and the reduced replication track lengths measured after UV in DNA fiber experiments can be interpreted as the result of successive uncoupling/recoupling cycles at each leading strand lesion encountered by a single RF during the labelling time. As WT cells also show a significant decrease of post-UV replication track lengths, we postulate that pol*η* does not prevent uncoupling but rather allows rapid recoupling of the stalled leading strand *in vivo*, as recently shown *in vitro*^42^.

On naked DNA templates, uncoupling can proceed to more than 1 kb past a leading strand CPD^18, 51^. However, our data indicate that uncoupling is limited in length *in vivo*, even when pol*η* is not functional, and rarely exceeds 450 nucleotides. As we could observe longer ssDNA after treatment with CPT using the same procedure^40^, this is unlikely to be due to DNA breaks during the DNA preparation, which could have led to the loss of longer ssDNA stretches. In absence of pol*η*, alternative tolerance pathways at the RF, including repriming and the recently described alternative bypass by polymerase theta (pol*θ*)^12^ should limit the extent of uncoupling. Uncoupling may also be restricted *in vivo* by topological constraints, chromatin ahead of the RF or checkpoint activation. In support of the latter hypothesis, Rad53 has been shown to restrict CMG helicase activity in the yeast purified replisome system^52^.

Numerous recombination-based mechanisms have been proposed to restart stalled RFs, such as HR or template switch. Moreover, fork reversal, which involved the key HR protein RAD51, was proposed as a universal response to replication stress in human cells^33^. We, however, did not observe any structural evidence of recombination or fork reversal in the early response to UV damage, even when pol*η* is not functional. This is in agreement with previous TEM studies, which showed that reversed forks are observed in response to bulky DNA lesions in yeast mutants inactivated for the checkpoint pathway^5^ or unable to efficiently reprime replication^53^. Accordingly, recent reports have unravelled a competition between fork repriming and fork reversal^54^ and distinct roles of fork remodelling and protection factors in response to HU or MMC^55^. It is therefore more than likely that the cell type and the origin of the replication stress dictate the choice between repriming and reversal. We further demonstrate here that on-the-fly direct bypass by pol*η*is the favoured early response to UV lesions in skin fibroblasts, outcompeting these two mechanisms.

### Repriming as a damage tolerance mechanism that compensates polη deficiency

Our structural data showed that repriming is the alternative mechanism used to promote RF progression when on-the-fly pol*η*-dependent TLS is not possible. We therefore propose that repriming constitutes a tolerance mechanism *per se*. This blockage-skipping mechanism was characterized in bacteria and, although not yet fully understood, also described in yeast and human cells^19, 25, 33, 56–58^.

Recently, in silico prediction based on conserved motifs with archea/eukaryotes AEP superfamily of primase identified a novel primase-polymerase, named PrimPol, in most eukaryotes, including mammals^20–22, 59^. We confirmed that PrimPol depletion further impaired RF progression in irradiated XPV cells and provided structural evidence that PrimPol can indeed reprime leading strand synthesis *in vivo*. However, our TEM analysis also suggests that its primase activity has a limited role after UV, even in a pol*η*-deficient background, which correlates with the modest impact of PrimPol deficiency on cell survival^45^ (data not shown) and the observation that PrimPol is rather involved in an adaptive response to multiple genotoxic exposures^54^. PrimPol also possesses a TLS polymerase activity. However, its active site is too constrained to accommodate UV lesions^60^, which it may rather skip through primer realignment^20, 22, 60, 61^. Therefore, PrimPol could contribute to RF progression in a primase-independent manner when pol*η* is not functional. Interestingly, we also identified a late function for PrimPol in the clearance of RPA accumulated on chromatin, suggesting a potential role in gap repair. Whether this involvement requires its primase and/or its lesion skipping activities remains unclear. PrimPol has been recently shown to be controlled by the ATR checkpoint^54^, a pathway that is over-activated in UV-exposed pol*η*-deficient cells^24^. It would therefore be interesting to assess the impact of PrimPol on UV-induced mutagenesis in XPV cells.

Beyond PrimPol, the polα primase-polymerase remains the only other currently identified candidate for repriming. The *E. coli* replisome possesses the inherent capacity to tolerate lesions in the leading strand by DnaG primase-dependent *de novo* priming^57^, a mechanism that competes with TLS^62^. Reconstitution of yeast replisome with purified proteins showed that repriming of a stalled leading strand by polα is rather inefficient^18^. Yeast cells do not possess a PrimPol homologue but TEM analysis revealed that repriming occurs *in vivo*^5, 53^. It is therefore most likely that accessory factors, post-translational modifications and/or RF remodelling processes promote polα (or a currently unidentified other primase) access to the stalled leading strand *in vivo*. Their identification would provide further mechanistic insights into the repriming mechanism in eukaryotes.

### Dual requirement for RAD51 at and behind RFs

We report two spatiotemporally and mechanistically distinct functions of RAD51 in response to UV damage.

Our data suggest that recombinational tolerance processes take place behind replication forks in a post-replicative manner for gap-filling in response to UV. Our TEM analysis leads to the original observation of recombination-related intermediates occurring *in vivo* on genomic DNA in human cells. Previous studies conducted in budding yeast evidenced that UV-induced recombination does not rely on double-strand breaks but on ssDNA gap processing^48^. Our *in vitro* studies showed that the five-branched structures observed *in vivo* are similar to intermediates formed during RAD51-dependent strand invasion processes, indicating that they can form independently of the presence of a replication fork and further supporting the idea that they form during post-replicative gap filling. Furthermore, their architectures are compatible with the models of template switch and recombinational gap repair described in literature^2^. Further work is needed to unravel the molecular mechanisms underlying their formation and resolution and determine if they are intermediates in the formation of UV-induced SCEs.

In addition to its well-documented role in HR, recent studies shed light on non-canonical roles of RAD51, to escort and stabilize ongoing RFs, but also to promote the restart of stalled RFs through their remodelling and the inhibition of their degradation by MRE11^26, 63–66^. Interestingly, it was reported that the strand-exchange activity of RAD51 is not required for fork protection^67^ and that RF restart and HR repair involve two different RAD51-mediated pathways^64^. Our data strongly suggest that RAD51 indeed works at stalled RFs in a strand-exchange-independent manner to sustain their progression when encountering UV lesions. However, contrary to previous reports^26, 33, 63^, we showed that RAD51 inhibition or depletion led to a decreased number of UV-specific post-replicative gaps. Hence, we hypothesize that RAD51 may contribute to repriming. As we observed that DNA2 inhibition partially restores post-UV track lengths, RAD51 may act after repriming occurred, by protecting the newly generated 5’ extremity from nucleolytic activity. Hence, our data suggests that reversed forks are not mandatory entry points for neo-synthesised DNA degradation, which can proceed from gap-delimiting dsDNA-ssDNA junctions. Interestingly, in contrast to what was observed after other genotoxic insults^40, 63, 65, 68, 69^, MRE11 does not trigger fork degradation after UV. This further supports the notion that the type of damage influences the choice of RI processing mechanisms^55^.

In addition to its role in RF protection, we cannot rule out that RAD51 may directly stimulate repriming. Yeast Pol*α* activity on the leading strand is inhibited by RPA *in vitro*^18^ and a direct interaction between RAD51 and pol*α* has already been reported in *Xenopus* extracts^69^. It is therefore tempting to speculate that RAD51 can promote access of pol*α* to the stalled leading strand. Similarly, although RPA was shown to recruit PrimPol on DNA, short ssDNA templates saturated by RPA are not good substrates for PrimPol primase activity^61, 70, 71^. Given that we showed that ssDNA generated at the RFs is limited in length, replacing RPA with RAD51 could also favour repriming by PrimPol. Further experiments are warranted to clarify how RAD51 mediates replication fork protection and restart.

### Alternative TLS polymerases and Mutagenesis in XPV syndrome

In WT cells, the bypass of 6,4-PPs is achieved through Polζ-dependent gap filling^25, 72^. Several studies demonstrated that polζ, polκ, and to a lesser extent polι, partially compensate pol*η* deficiency in XPV cells, leading to mutagenic CPD bypass^9, 10, 12^. Here, we showed that polζ and polκ do not contribute to the progression of RFs even in the absence of pol*η*, indicating that alternative TLS of CPDs proceeds post-replicatively. Importantly, in budding yeast, the mismatch repair (MMR) pathway operates coupled to DNA replication^73^ and is therefore less efficient on mismatches created during polζ-mediated gap filling^74, 75^. Hence, the inefficiency of mismatch correction caused by the delayed bypass of TT-CPD imposed by pol*η* deficiency may further amplify the intrinsic error-proneness of the alternative pathways taking place in XPV cells.

A recent study has identified polθ as the inserter TLS polymerase initiating the mutagenic bypass of CPDs before further extension by polζ or polκ^12^. Intriguingly, polθ deficiency leads to impaired RF progression after UV, in a synergic manner with pol*η* deficiency^12^. This raises the possibility that the insertion made by polθ at the RF stimulates repriming, a hypothesis that could be tested in future TEM experiments. It would also be important to understand why alternative TLS polymerases do not bypass UV lesions at the RF, a situation that may favour MMR-mediated correction of their errors. We have previously shown that SUMOylation of pol*η* promotes its association with the replication machinery^38^. The SUMOylation site of pol*η*is not conserved in other Y family DNA polymerases, suggesting a regulation specific to pol*η*. Hence, TLS polymerases could be differentially regulated for access of the vicinity of RFs. Alternatively, when pol*η* is not functional, the slower efficiency and/or kinetics of photoproduct bypass may also leave time for repriming to occur and RF to progress before TLS is completed.

Another open question concerns the respective contributions of TLS and HR to gap repair. Previous studies in budding yeast proposed a redundancy between TLS and HR to process UV-induced post-replicative gaps^48^. In mammalian cells, depletion of pol*η* and/or polθ leads to a higher number of RAD51-dependent SCEs after UV^12^, suggesting that HR compensates for TLS deficiency. Elsewhere, a competition between TLS and HR pathways has been extensively suggested^47, 76–79^. The suggested role of TLS polymerases in HR itself further blurry the picture^80–82^. Here we show that RAD51 is preferentially recruited behind RFs and that its inhibition sensitizes cells to UV, both in pol*η*-deficient and pol*η*-proficient cells. One possibility could be that the whole post-replicative tolerance is mediated through HR events but that they involved synthesis steps made by TLS polymerases. On the other hand, as the recombination intermediates we observed are rare compared to replication forks, RAD51 could also have a HR-independent function behind RFs, similar to its role at RFs, through protection of the gaps in order to allow their fill in by canonical TLS.

Altogether, our data allow drawing a chronology of DNA damage tolerance mechanisms in response to UV lesions with pol*η*acting first at the RF, followed by repriming supported by a fork protection function of RAD51 if pol*η* is not efficient and further post-replicative gap filling through TLS and/or HR. Hence, our study supports a collaboration between the different DDT pathways for replication completion of damaged DNA and the idea that the DDT pathway choice is dictated by the type of lesion and the most efficient mechanism available, giving a priority to the continuity of DNA synthesis.

## Aknowledgements

The authors are indebted to L. Blanco for the kind gift of anti-PrimPol antibody. The authors thank L. Blanco, A. Doherty, G. Mazon and all members of the P.L.K. lab for helpful discussions and J.H. Guervilly for critical reading of the manuscript. The authors thank Barbara Ben-Yamin and Raphaël Corre for the RT-qPCR experiments. This work was supported by grants from La Ligue Nationale contre le Cancer “Equipe labellisée”, INCa (INCa-PLBio2016-159), INCa-DGOS-Inserm 12551, ANR (FIRE 17-CE12-0015) and Paris-Sud University (MRM). E.D. was supported by INCa (PLBio2016-144). Y.B. was supported by the SIRIC SOCRATE, Gustave Roussy (INCa-DGOS-Inserm 12551).

## Authors contributions

Y.B. performed cell fractionation experiments, TEM experiments and cell survival assays. C.P. performed the analysis of PrimPol recruitment by cell fractionation. E.M.T. and P.D. performed the *in vitro* strand exchange assay. E.D. performed iPOND experiments, IF and PLA experiments and DNA fiber assays. Y.B., P.L.K., E.L.C. and E.D. designed the study and analysed the results. Y.B. and E.D. wrote the manuscript with the help of P.L.K. and E.L.C.

## Material and Methods

### Cell culture and treatments

SV40-transformed normal (MRC5-V1) and XPV (XP30RO) human fibroblasts were grown in Minimum Eagle Medium (MEM, Gibco) supplemented with 10% fetal calf serum, 100 U/μl penicillin and 100 μg/ml streptomycin under a 5% CO_2_ atmosphere. XP30RO cells stably complemented by wild-type pol*η* (XPV^polη^ cells) or pol*η* mutants were described elsewhere^83, 84^ and were grown in the presence of 100 μg/ml zeocin (Invivogen). Mouse embryonic fibroblasts (kind gift from R. Wood) were cultured in Dulbecco’s Modified Eagle Medium (DMEM, Gibco) supplemented with 10% fetal calf serum, 100 U/μl penicillin and 100 μg/ml streptomycin under a 5% CO_2_ atmosphere. For UVC irradiation (254 nm), cells were washed with pre-heated Phosphate Buffered Saline (PBS) and irradiated without any medium at a fluency of 0.65 J.m^-2^.s^-1^. Stock solutions of B02, mirin (Sigma) and DNA2i (MedChemExpress) were prepared in DMSO.

### siRNA transfection

siRNAs purchased from Dharmacon were used to transiently down-regulate the expression of *POLH* (5’-GAAGUUAUGUCCAGAUCUU-3’), *POLK* (On-TARGET plus SMART pool), *PRIMPOL* (On-TARGET plus SMART pool), *RAD51* (On-TARGET plus SMART pool, siGENOME duplex 2) and *REV3L* (On-TARGET plus SMART pool). Unspecific siRNAs (siNT) were used as control. Cells were transfected with 30 nM of siRNAs using Interferin (PolyPlus) according to manufacturer’s instructions, and were treated 48 h to 72 h after transfection. For RAD51 depletion, 15 nM of siRNAs were used for 36 h.

### Proliferation assay

Cells were plated at 1.5 x 10^5^ per well of a six-well plate 24 h before UVC exposure. Cells were treated as indicated, incubated for 72 h and then counted with trypan blue staining using a Neubauer haemocytometer.

### Cellular fractionation

Cells were harvested by trypsinization, washed in PBS and then subjected to cell fractionation. Cells (0.5-1.10^6^) were resuspended in 1 ml of CytoSKeleton (CSK) 100 buffer (100 mM NaCl, 300 mM Sucrose, 3 mM MgCl_2_, 10 mM PIPES pH 6.8, 1 mM EthyleneGlycol-tetra-acetic Acid (EGTA), 0.2 % Triton X-100, and protease inhibitor cocktail (Roche)) and incubated for 15 min on ice. Samples were centrifuged at 7000 rpm for 5 min at 4°C, and washed once in CSK buffer. The pellet, corresponding to the insoluble protein fraction, was solubilized in 100 μl lysis buffer (10 mM Tris-HCl pH 7.5, 20 mM NaCl, 0.4 % SDS, protease inhibitor cocktail) supplemented with 150 U/ml benzonase (Millipore) for 30 min at 37°C. For whole-cell extracts (WCE), cell pellets were directly lysed in lysis buffer with benzonase for 10 min at room temperature. Protein amounts present in samples were evaluated using Nanodrop (Thermo) or Bradford assay. Proteins were denaturated for 10 min at 90°C in the presence of Laemmli buffer.

### iPOND

Cells were irradiated with 25 J.m^-2^ UVC and pulse-labelled with 10 μM EdU (Invitrogen) for 10 min. In Supplementary Figure 4d, the UVC dose was reduced to 5 J.m^-2^ and the labelling period was extended to 20 min. iPOND was performed immediately after the pulse or after a chase in fresh medium supplemented with 10 μM thymidine (Sigma) as already described^38^. Briefly, cells were crosslinked with 1% formaldehyde (Sigma), harvested by scrapping and permeabilized in PBS with 0.5% TritonX100. Biotin-azide (Molecular Probes) was conjugated to EdU by click chemistry for 2 h in click reaction buffer (10 mM sodium-L-ascorbate, 10 mM biotin-azide, 2 mM CuSO4 in PBS). Cells were lysed in iPOND lysis buffer (10 mM Hepes-NaOH pH 7.9, 100 mM NaCl, 2 mM EDTA, 1 mM EGTA, 1 mM PMSF, 0.2% SDS, 0.1% sarkozyl, antiproteases) and sonicated on a Bioruptor device. Solubilized chromatin was retrieved by centrifugation and subjected to streptavidin pull-down overnight (Dynabeabs MyOne Streptavidine C1, Invitrogen). After bead washing, pull-downed proteins were eluted by boiling in 1x Laemmli buffer at 95 °C for 30 min.

### Western Blot

Samples were subjected to SDS-PAGE followed by PVDF membrane transfer. Membranes were immunoblotted with the following antibodies: mouse anti-β-actin (AC-15, Sigma #A5441), mouse anti-β-catenin (cl14, BD Biosciences, #610153), rabbit anti-histone H2B (V119, Cell Signaling #8135), rabbit anti-histone H3 (Abcam #ab1791), mouse anti-LaminA/C (Santa Cruz #sc-7292), mouse anti-PCNA (PC10, Santa Cruz #sc-56), goat anti-POLA1 (G-16, Santa Cruz #sc-5921), goat anti-POLD1 (C-20, Santa Cruz #sc-8797), rabbit anti-polκ (Bethyl #A301-975A), rabbit anti-pol*η* (Abcam #ab17725, Bethyl #A301-231A), mouse anti-pol*η* (B-7, Santa Cruz #sc-17770), rabbit-anti PrimPol (kind gift from L. Blanco), rabbit anti-RAD51 (Calbiochem #PC130-100UL), mouse anti-RPA32 (Calbiochem #NA19L) and mouse anti-RPA70 (Calbiochem #NA13).

### Genomic DNA preparation and RIs enrichment

Replication intermediates were isolated as follows. Cells subjected to the indicated treatments were washed with PBS and DNA was crosslinked by two rounds of incubation on ice in a dark room for 5 min with 10 μg/ml trioxalen (Sigma) followed by irradiation with 45 kJ UVA, in order to generate interstrand crosslinks in the genomic DNA. Cells were then washed and harvested by scraping, then subjected to nuclei extraction (Nuclei EZ prep kit, Sigma). Genomic DNA was isolated using DNAzoL reagent (Invitrogen) following manufacturer’s instructions and resuspended in water. Accurate deproteinization, which is necessary to avoid intra- and inter-molecular crossovers (Benureau, in preparation), was performed by two rounds of hydrolysis by 1 mg/ml proteinase K overnight at 55°C in the presence of 1% SDS, followed by a gentle phenol/chloroform extraction. After precipitation, genomic DNA was treated at 37°C for 2 h with RNaseA (DNase and protease-free,Thermo Scientific) and BamHI (New England Biolabs).

Solution was adjusted to 300 mM NaCl and loaded onto a benzoylated naphtoylated DEAE-cellulose (BND, Sigma) column chromatography. Fractions corresponding to dsDNA molecules were collected with DS Elution buffer (10 mM Tris-HCl pH 7.5, 5 mM EDTA and 0.8 M NaCl). Genomic DNA fragments containing ssDNA regions were eluted by the addition of SS Elution buffer (10 mM Tris-HCl pH 7.5, 5 mM EDTA and 1 M NaCl, 0.9% caffeine (Sigma)). SS fractions were pooled, precipitated with isopropanol, resuspended in water, quality-controlled by agarose gel electrophoresis and dosed with Nanodrop. In order to ensure that RIs structures are not lost during the whole procedure, the UVA/trioxalen crosslinking efficiency was systematically checked by TEM observation after heat/formamide denaturation of the samples.

### Electron Microscopy

We specifically developed a method to optimize recruitment, deployment of long ds-ssDNA molecules and their visualization in darkfield imaging mode. We have combined BAC (Benzyl-dimethyl-alkyl-ammonium chloride) hyperphase method with positive staining direct adsorption method. The hyperphase method initially developed by Vollenweider et al.^85^ allows to stretch DNA intermediates and to avoid secondary structures and random intra- or inter-molecular crossovers. The direct adsorption method consists in the treatment of the carbon film with a glow discharge in presence of amylamine (positive charges) as initially demonstrated by Dubochet et al.^86^. It favours the recruitment of DNA intermediates onto the surface, allows a positive staining of DNA by uranyl acetate and its observation in dark-field imaging mode. We obtain high-resolution images with an optimal contrast^40^.

Briefly, ssDNA-enriched samples were added to a mix of formamide, BAC and glyoxal, to generate a hyperphase film onto the surface of a water bath. The DNA was picked up on a positive functionalized carbon-coated grids by a brief contact with the hyperphase film surface. Grids were then washed in a large volume of purified milliQ water followed by 100% ethanol and dried with filter paper. Finally, molecules adsorbed onto the grids were positively stained with 2% uranyl acetate and dried. Samples were finally visualized in annular dark field mode, filtered in zero loss, with a Zeiss EM902 transmission electron microscope equipped with an annular Electron Energy Loss Spectrometer (EELS). Electron micrographs were acquired using a Veletta high-resolution CCD camera associated to a dedicated iTEM software (Olympus, Soft Imaging Solutions) with a 30,000 X magnification. Inlays correspond to a 140,000 X magnification.

### Strand exchange assay

Rad51 and RPA proteins from yeast *Saccharomyces cerevisiae* were purified as previously described^87^. 13 μM Rad51 were incubated with 40 μM of circular viral ssDNA (+) PhiX174 (previously purified in ion exchange MiniQ column) for 3 minutes, in 10 mM Tris-HCl pH 7.5, 100 mM NaCl, 3 mM MgCl_2_, 2 mM ATP, 1 mM DTT and 2 mM Spermidine, and 1.3 μM RPA was then added into reaction for 10 minutes. 20 μM of linear dsDNA PhiX174 replicative form I (Phix174 RFI homologous plasmid purchased from New England Biolabs and linearized by digestion with the restriction enzyme XhoI or PstI followed by phenol-chloroform extraction) was added into the reaction for 30 minutes, to a 10 μL final volume. The reaction was performed at 37°C. Reactions were crosslinked with psoralen (final concentration 10 μg/mL) during 4 min under UVA (crosslink density of 1 per 200-300 nt) and stopped using 0.5 mg/mL Proteinase K, 1% SDS, 12.5 mM EDTA and incubated overnight at room temperature. For TEM analysis, 4 strand exchange reactions were pooled, DNA was purified using phenol-chloroform and ethanol precipitation then spread using BAC hypophase method (as for *in vivo* RIs TEM samples preparation).

### DNA fiber assay

Cells were incubated for 20 min in medium supplemented with 25 μM iododeoxyuridine (IdU, Sigma). Cells were washed, irradiated with UVC and further incubated in medium supplemented with 50 μM chlorodeoxyuridine (CldU, Sigma) for 20 to 60 min as indicated. Alternatively, cells were first labelled with 25 μM CldU, then with 25 μM IdU. 1,000 cells, resuspended in 2 μl of PBS, were spotted on microscope glass slides and lysed in 7 μl of spreading buffer (200 mM Tris-HCl pH 7.5, 50 mM EDTA pH 8.0, 10% SDS). DNA was spread by tilting the slides. DNA was fixed in methanol/acetic acid 3:1 and denatured in 2.5 M HCl for 1 h at room temperature. Samples were blocked in blocking buffer (1% BSA, 0.1% Tween20 in PBS) for 1 h. IdU, CldU and total DNA were detected using the following protocol^88^: (1) mouse anti-BrdU (Becton Dickinson, 1/20, IdU) + rat anti-BrdU (Abcys, 1/100, CldU) for 1 h at room temperature, (2) goat anti-mouse AF594 + goat anti-rat AF488 for 30 min at 37°C, (3) mouse anti-single stranded DNA (Millipore, 1/100) for 45 min at 37°C, (4) rabbit anti-mouse AF350 for 20 min at 37°C and (5) goat anti-rabbit AF350 for 20 min at 37°C. All fluorescent antibodies were from Molecular Probes and used at 1/100 dilution. Antibodies were diluted in blocking buffer. Samples were washed four times in PBS between each antibody incubation. Samples were mounted in fluorescent mounting medium (DAKO). Images were acquired and analysed on an Axio Imager Z1 microscope using the Axio Vision software (Zeiss). Broken replication tracks were excluded from the analysis. Statistical significance was assessed by the non-parametrical Mann-Whitney test.

### Immunofluorescence

Cells were pre-extracted in CSK100 buffer for 5min on ice under gentle agitation. Cells were fixed in 4% paraformaldehyde for 20 min and permeabilized in methanol at −20 °C for 10 s. Cells were incubated for 1 h at room temperature with primary antibodies (rabbit anti-RAD51 Calbiochem PC130 1/200 and mouse anti-PCNA Santa Cruz PC10 1/500) diluted in IF buffer (3% BSA, 0.5% Tween 20 in PBS). Cells were washed three times in PBS and stained for 30 min with secondary antibodies from Molecular Probes (goat anti-rabbit AF488 1/1,000, goat anti-mouse AF594 1/1,000). Coverslips were mounted in fluorescent mounting medium (DAKO) supplemented with DAPI (1.5 μg/ml). Images were acquired on an AxioImager Z1 microscope using the Axio Vision software (Zeiss). Intensity was quantified with ImageJ software.

### In situ Proximity ligation assay

PLA to nascent DNA was performed as already described^38^. Briefly, cells were irradiated with UVC (25 J.m^-2^) and incubated for 30 min. 10 μM EdU was added for the last 7 min of the incubation. Cells were pre-extracted in CSK100 buffer and fixed prior to biotin-azide conjugation to EdU. Cells were incubated with primary antibodies against RAD51 (rabbit anti-Rad51 1/200, Calbiochem PC130) and biotin (mouse anti-biotin 1/6,000, Jackson ImmunoResearch #200-002-211). PLA and EdU counterstaining were performed according to the manufacturer’s instructions using the Duolink In Situ Red kit (Sigma) and goat anti-mouse Alexa Fluor 488 antibody.

### RT-QPCR

Total RNA extraction was performed using Maxwell RSC simply RNA cell kit (Promega) according to the manufacturer’s instructions. 1 μg RNA was reverse transcribed to cDNA with Maxima H minus Reverse Transcriptase (Thermo Scientific) using the following amplification program: 10 min at 25°C, 60 min at 50°C, 5 min at 85°C. Samples were treated with RNaseH (Thermo Scientific) for 20 min at 37°C and 10 min at 65°C. qPCR was performed with SYBR Green ROX qPCR Master Mix on CFX96 RealTime Systen (Biorad) using the following primers: hREV3L-F: ACCCCCGGAGTACCACTTATCCAGC, hREV3L-R: CCGGAGATATGGTGCCTTGGACA, hGAPDH-F: ACCACAGTCCATGCCATCAC, hGAPDH-R: TCCACCACCCTGTTGCTGTA.

---

**Supplementary Figure 1.**
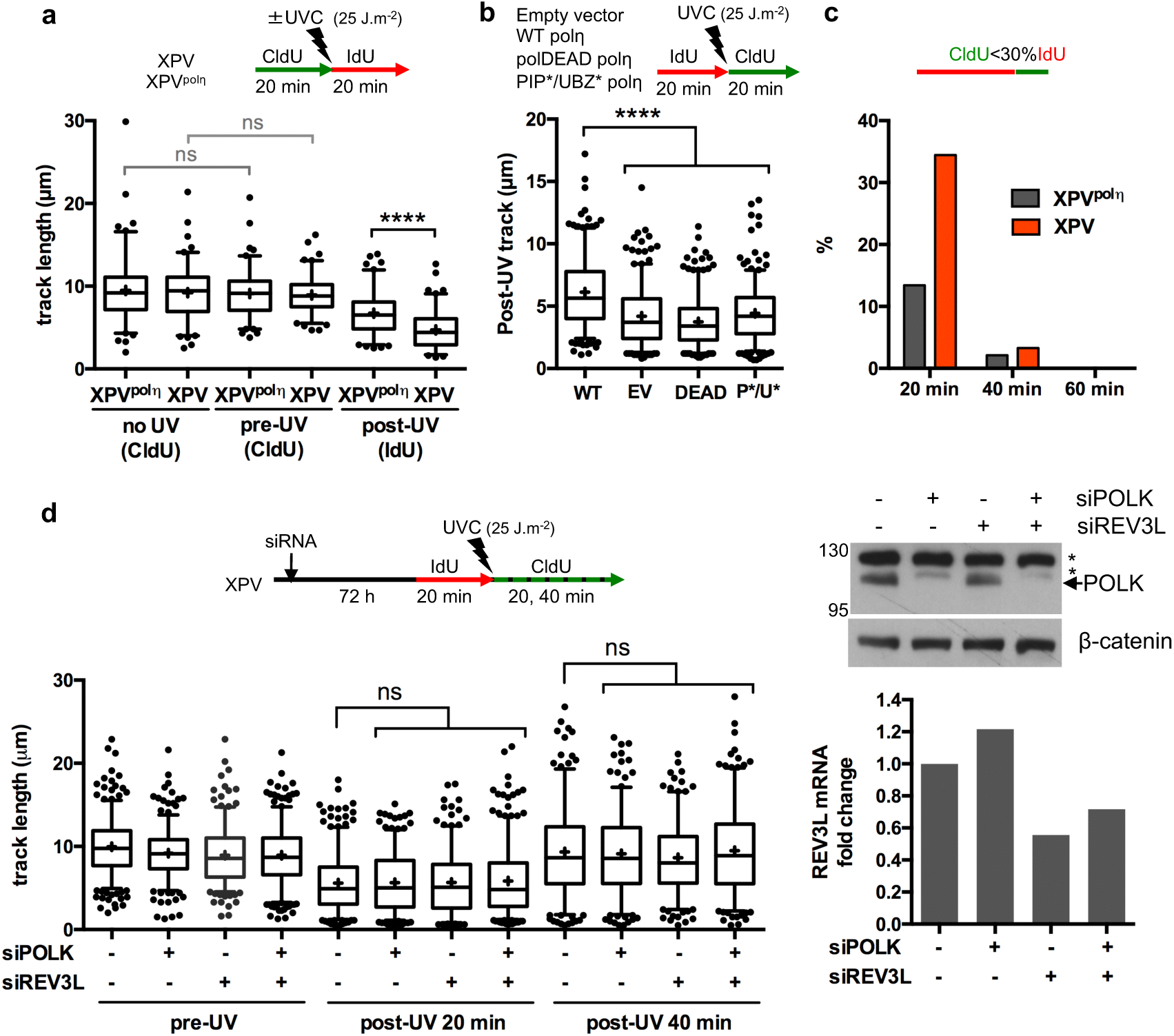
Polη, but not polκ or polζ, promotes replication fork progression after UV (supporting data for Figure 1). (**a**) Cells were first pulse-labelled with CldU (20 min), irradiated at 25 J.m^-2^ and pulse-labelled with IdU (20 min). Replication fork progression was analysed by DNA fiber assay (n=100). (**b**) XPV cells reconstituted with wild-type pol*η* (WT), catalytically inactive pol*η* (polDEAD, DEAD), pol*η* mutated in its PCNA- and ubiquitin-interacting motifs (PIP*/UBZ*, P*/U*) or an empty vector (EV) were treated as in Figure 1b. Distribution of post-UV track lengths (CldU) is shown (n=300). (**c**) The proportions of replication signals with CldU length shorter than 30% of the corresponding IdU length were calculated for the experiment shown in Figure 1d. (**d**) XPV cells were depleted for POLK and/or REV3L, the catalytic sub-unit of pol*ζ*. Pre- and post-UV replication track lengths were measured on more than 240 signals. Efficiency of POLK and REV3L depletions were analysed by western blot and by RT-qPCR, respectively. Statistical analysis was made with the Mann-Whitney test (ns: not significant, *: P<0.05, ****: P<0.0001).

**Supplementary Figure 2.**
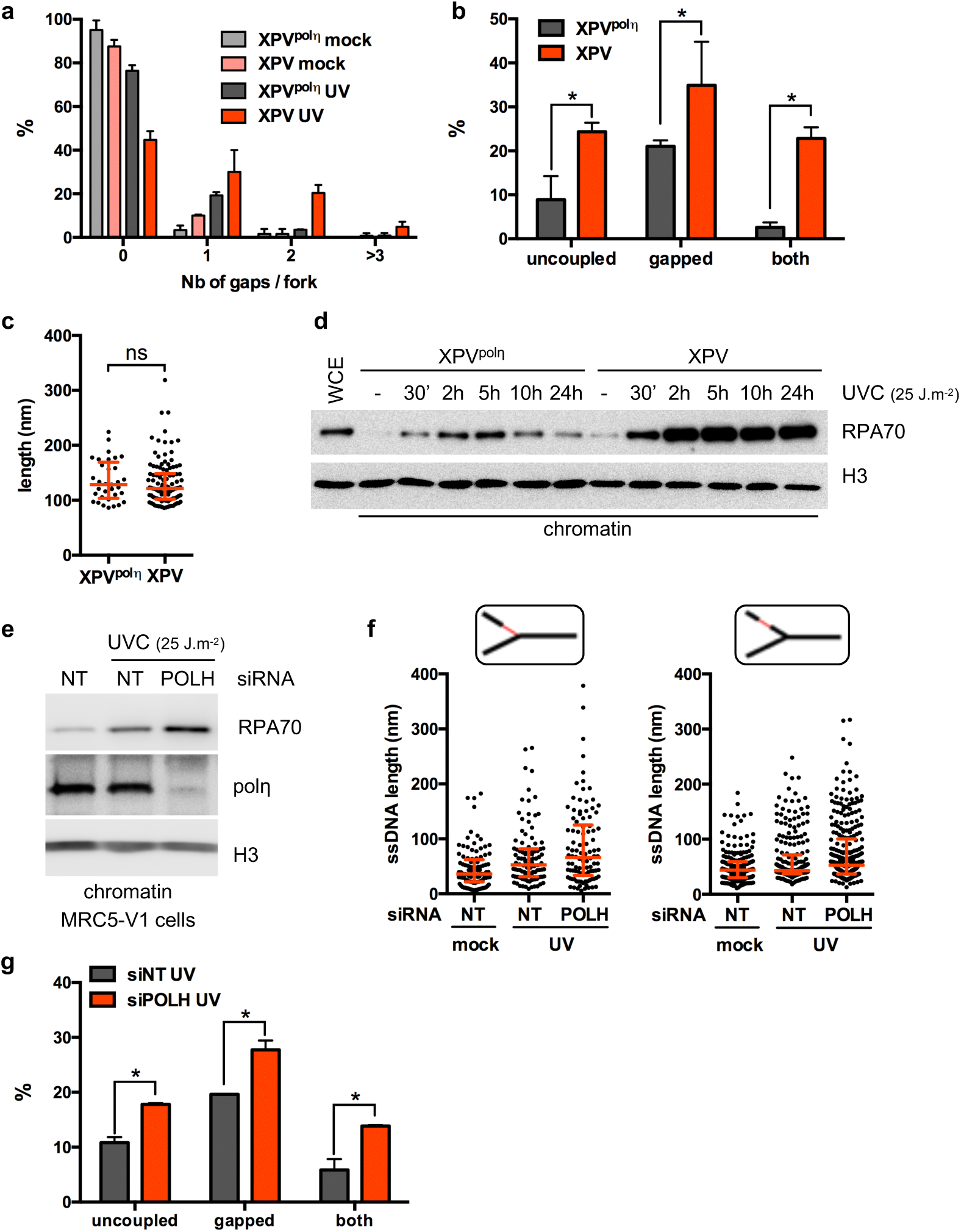
Polη deficiency leads to accumulation of uncoupled forks and post-replicative gaps after UV (supporting data for Figure 3). (**a**) Frequency distribution of the number of gaps per fork in XPV cells compared to XPV^polη^ cells (mean ± SD of two independent experiments). (**b**) Percentage of the analysed forks displaying only uncoupling, only post-replicative gap(s) or both features after UVC (mean ± SD of two independent experiments, multiple t-test, *: P<0.05). (**c**) Distribution of the lengths of the post-replicative gaps induced by UVC exposure (i.e. above the threshold defined in the text) in XPV and XPV^polη^ cells (pool of the two experiments, Mann-Whitney test, ns: not significant). (**d**) Time-point analysis of chromatin-bound RPA70 after UVC irradiation (25 J.m^-2^) in XPV^polη^ and XPV cells. Histone H3 was used as a loading control. WCE: whole-cell extract. (**e**) Chromatin-bound RPA70 was analysed 30 min after UVC (25 J.m^-2^) in MRC5-V1 cells following down-regulation of endogenous pol*η* by siRNA. (**f**) Distributions of the ssDNA lengths measured at (left panel) and behind (right panel) the elongation point in MRC5-V1 cells depleted for pol*η* (pool of two independent experiments). The 95^th^ percentiles of the distribution in untreated control cells were 106.3 nm and 108.3 nm at and behind RFs, respectively, and used as thresholds to calculate the percentage of uncoupled and gapped forks shown in Figures 3e,f. (**h**) Percentage of the analysed forks displaying only uncoupling, only post-replicative gap(s) or both features in MRC5-V1 cells exposed to UVC (mean ± SD of the two independent experiments, multiple t-test, *: P<0.05).

**Supplementary Figure 3:**
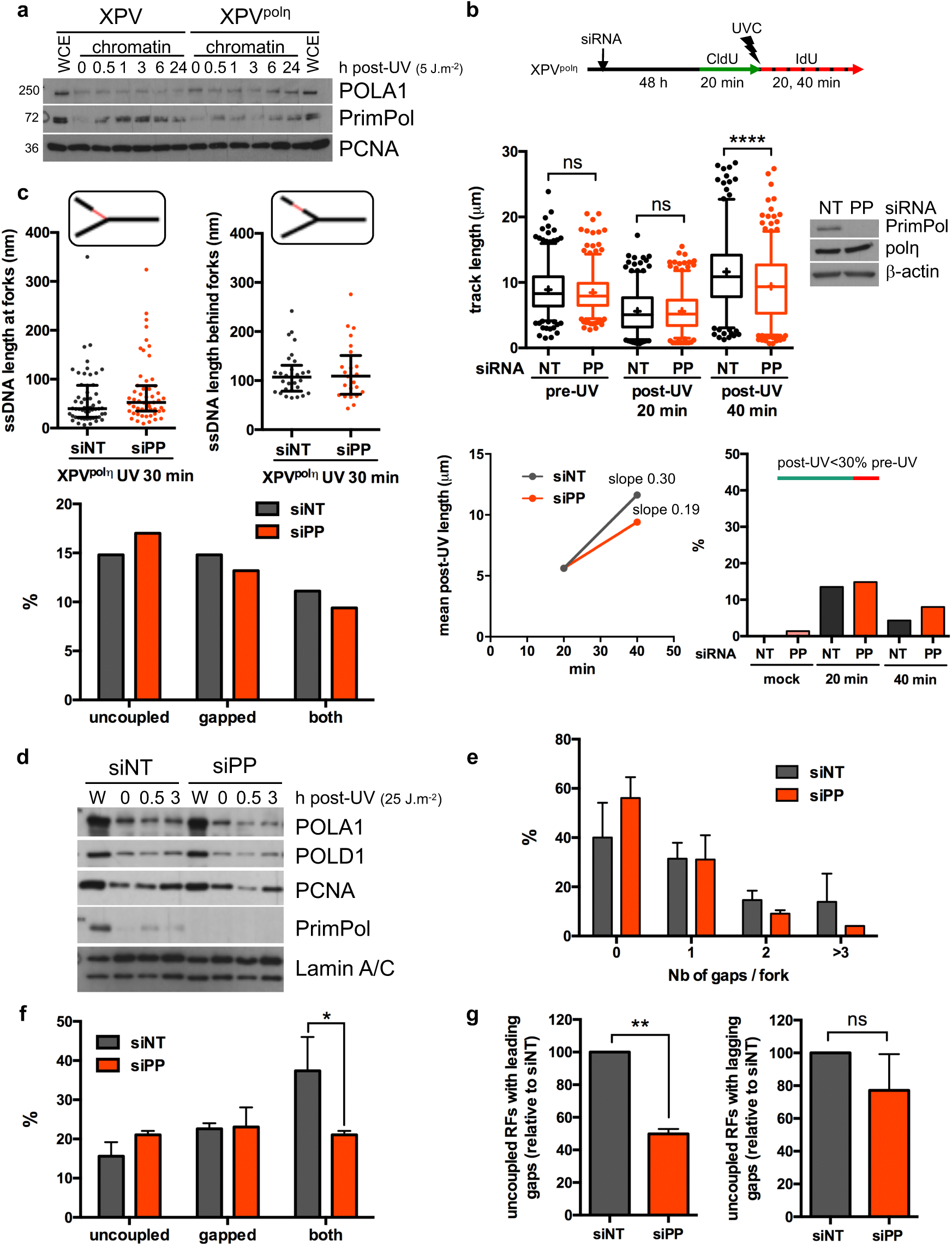
Impact of PrimPol depletion on RF progression and architecture after UVC (supporting data for Figure 4). (**a**) Cellular fractionation was performed at several time points following UVC irradiation at 5 J.m^-2^ in XPV and XPV^polη^ cells. Whole cell extracts (WCE) of unirradiated cells and chromatin-bound proteins were analysed by western blot using the indicated antibodies. (**b**) DNA fiber assay was performed in XPV^polη^ depleted for PrimPol (PP). siPRIMPOL efficiency was confirmed by western blot using the indicated antibodies. Upper panel show the distributions of replication track length (n>250, Mann-Whitney test, ns: not significant; ****: P<0.0001). Lower left panel shows the average post-UV length. Lower right panel show the proportion of replication signals with post-UV length shorter than 30% of the corresponding pre-UV length. (**c**) TEM analysis of PrimPol-depleted XPV^polη^ cells 30 min after UVC (25 J.m^-2^). The distributions of ssDNA track lengths measured at and behind the elongation point are shown in the upper left and upper right panel, respectively (median with interquartile range). Lower panel shows the proportion of RFs displaying only uncoupling, only post-replicative gap(s) or both features. (**d**) Fractionation of mock- and PrimPol-depleted XPV cells after 25 J.m^-2^. WCE (W) and insoluble protein fractions were analysed by western blot with the indicated antibodies. (**e**) Frequency distribution of the number of gaps per fork in mock- and PrimPol-depleted XPV cells (mean ± SD of two independent experiments). (**f**) Percentage of the RFs displaying only uncoupling, only post-replicative gap(s) or both features after UVC in mock- and PrimPol-depleted XPV cells (mean ± SD of two independent experiments, multiple t-test, *: P<0.05). (**g**) Relative number of uncoupled RFs with gaps located on the leading (left panel) or on the lagging (right panel) strand (mean ± SD of two independent experiments, t-test, ns: not significant, **: P<0.01).

**Supplementary Figure 4.**
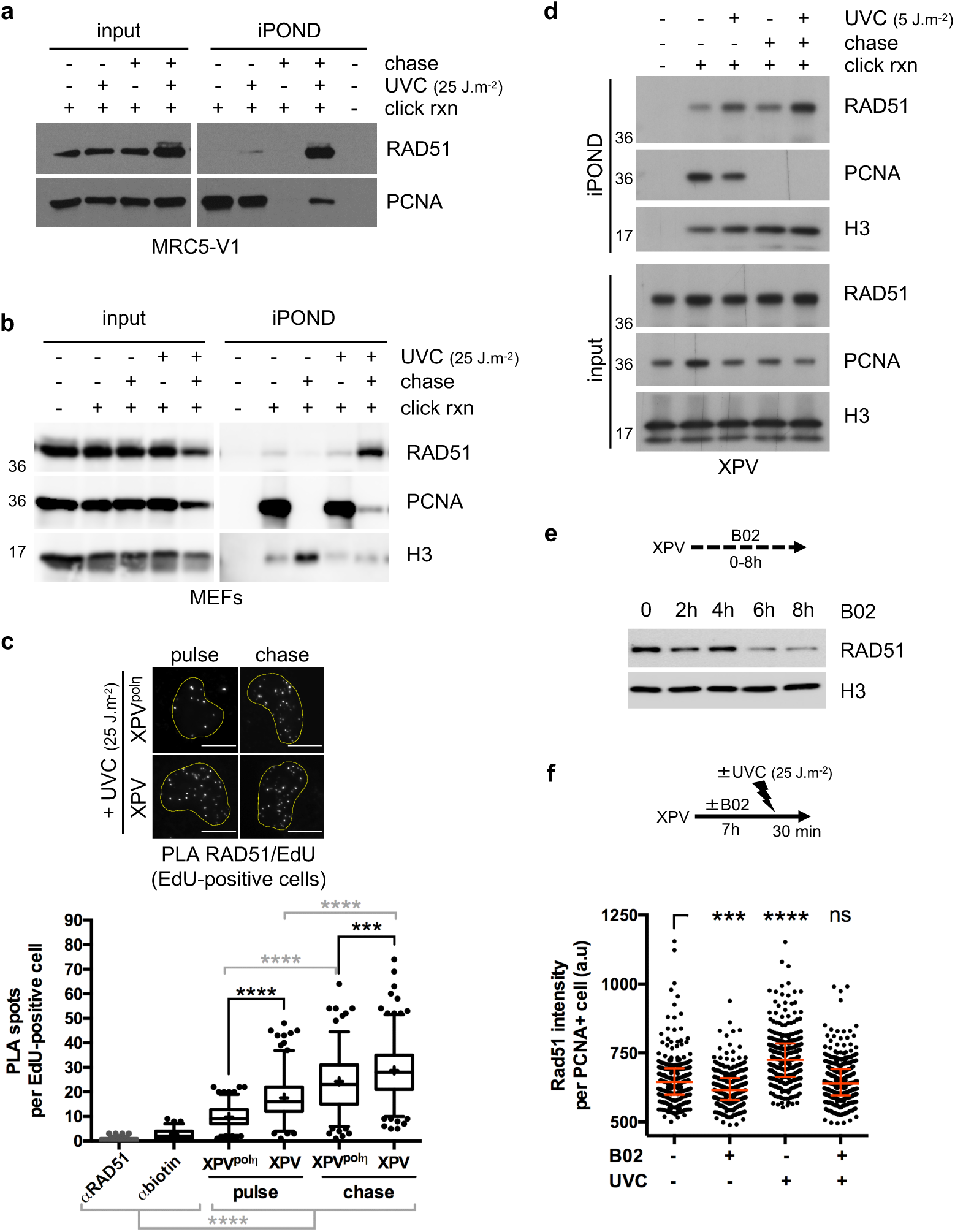
Dynamics of RAD51 at and behind replication forks after UV (Supporting data for Figure 5). (**a**) iPOND in MRC5-V1 cells. (**b**) iPOND in mouse embryonic fibroblasts (MEFs). Cells were irradiated at 25 J.m^-2^, pulse-labelled for 10 min with EdU and chased for 1 h in thymidine. (**c**) PLA between nascent DNA and RAD51 in XPV and XPV^polη^ cells after UVC. Cells were irradiated at 25 J.m^-2^ and incubated for 30 min with an EdU pulse performed during the last 7 min. When indicated, EdU was chased with thymidine for 1 h. Cells were pre-extracted and fixed before conjugation of biotin on EdU by click chemistry. PLA between EdU (biotin) and RAD51 was performed prior to EdU counterstaining. PLA reactions omitting one of the two primary antibodies were used as negative controls. The number of PLA spots was counted in EdU-positive cells, except in the RAD51 Ab only negative control in which spots were counted in random nuclei, and displayed as box-plots with 5-95 percentile whiskers (+: mean of the distribution, dots: outliers, Mann-Whitney test, ***: P<0.001, ****: P<0.0001). scale bar = 10 μm. (**d**) XPV cells were irradiated at 5 J.m^-2^, pulse-labelled for 20 min with EdU, chased for 1 h in thymidine before performing iPOND. (**e**) XPV cells were treated with 30 μM B02 for the indicated times. Chromatin-bound RAD51 amounts were assessed by cell fractionation. (**f**) XPV cells were treated with 20 μM B02 for 7 h prior to UVC irradiation (25 J.m^-2^). Cells were pre-extracted and fixed 30 min after irradiation. The amount of chromatin-bound RAD51 was assessed in S phase cells (PCNA-positive cells) by immunofluorescence (dot plot with median and interquartile range, Kruskal-Wallis test, ns: not significant, ***:P<0.001, ****: P<0.0001).

**Supplementary Figure 5.**
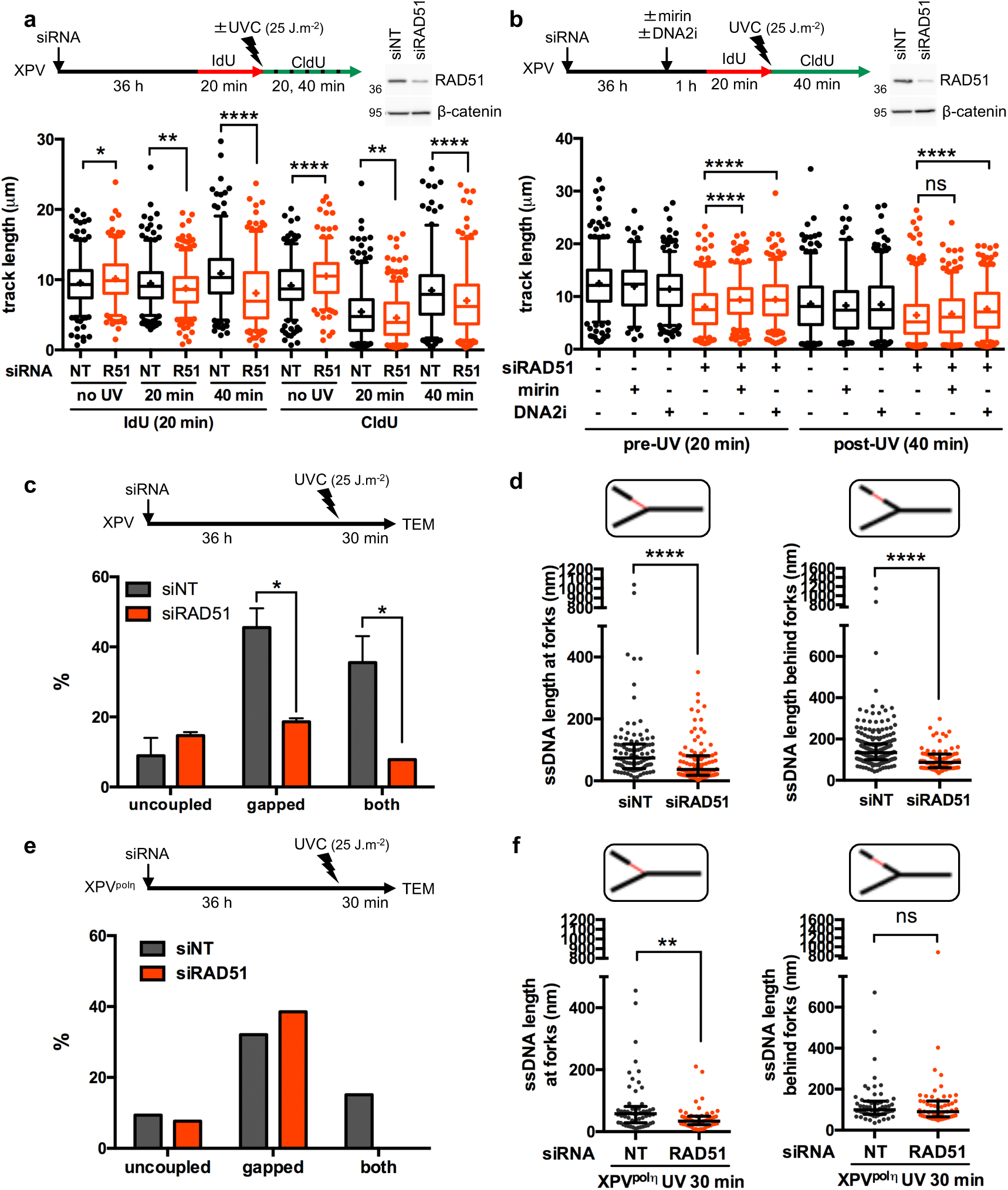
Depletion of RAD51 impairs replication fork progression and gap formation after UV (Supporting data for Figure 5). (**a**) XPV cells were depleted for RAD51 (R51) and DNA fiber assay was performed as described in Figure 1d (n> 250, Mann-Whitney test, ns: not significant, *: P<0.05, **: P<0.01, ***: P<0.001, ****: P<0.0001). (**b**) XPV cells were transfected with another siRNA targeting RAD51 and pre-treated for 1 h with 50 μM Mirin or 25 μM DNA2i. DNA fiber assay was performed as in Figure 5c (n>120). (**c**) The percentages of RFs displaying only uncoupling, only post-replicative gap(s) or both features 30 min after UVC (25 J.m^-2^) in mock- and RAD51-depleted XPV cells were determined by TEM (mean ± SD of two independent experiments, at least 50 RIs were analysed per condition in each experiment, multiple t-test, *: P<0.05). (**d**) The distributions of ssDNA lengths at and behind RFs are shown as dot plots with median and interquartile range (pool of two independent experiments, Mann-Whitney test, ****: P<0.0001). (**e**) As in c in XPV^pol^*^η^* cells (1 experiment, n>50). (**f**) As in d in XPV^pol^*^η^* cells (Mann-Whitney test, ns: not significant, **: P<0.01).

**Supplementary Figure 6.**
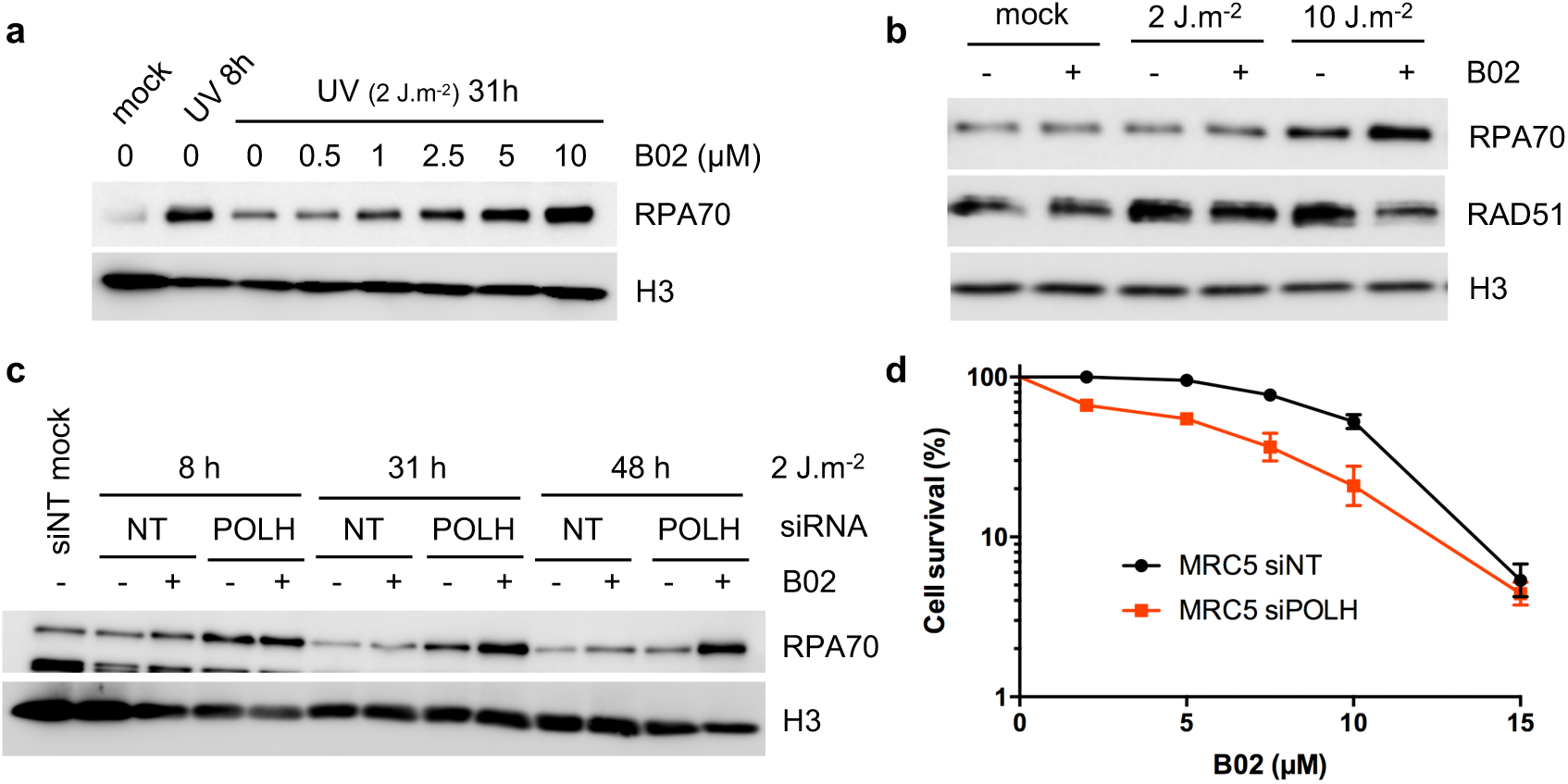
Long-term consequences of RAD51 inhibition in UV-irradiated cells (Supporting data for Figure 6). (**a**) Dose-dependent effect of B02 on chromatin-bound RPA in XPV cells after UVC irradiation (2 J.m^-2^). (**b**) XPV^polη^ cells were irradiated with UVC at the indicated doses and treated or not with 7.5 μM B02. Cellular fractionation was performed 24 h after irradiation. (**c**) MRC5-V1 cells were depleted for pol*η* and irradiated with UVC (2 J.m^-2^). When indicated, 10 μM B02 was added after irradiation. Chromatin-bound RPA was analysed by cellular fractionation. (**d**) Cell survival after 2 J.m^-2^ in MRC5-V1 cells depleted or not for pol*η* and treated with increasing doses of B02 as indicated. 100% represents the number of cells after 72 h in the DMSO condition for each siRNA. Values are the mean ± SD of three independent experiments.

